# Lecithin:Retinol Acyl Transferase (LRAT) induces the formation of lipid droplets

**DOI:** 10.1101/733931

**Authors:** Martijn R. Molenaar, Tsjerk A. Wassenaar, Kamlesh K. Yadav, Alexandre Toulmay, Muriel C. Mari, Lucie Caillon, Aymeric Chorlay, Maya W. Haaker, Richard W. Wubbolts, Martin Houweling, A. Bas Vaandrager, Fulvio Reggiori, Abdou Rachid Thiam, William A. Prinz, J. Bernd Helms

## Abstract

Lipid droplets are unique and nearly ubiquitous organelles that store neutral lipids in a hydrophobic core, surrounded by a monolayer of phospholipids. The primary neutral lipids are triacylglycerols and steryl esters. It is not known whether other classes of neutral lipids can form lipid droplets by themselves. Here we show that production of retinyl esters by lecithin:retinol acyl transferase (LRAT) in yeast cells, incapable of producing triacylglycerols and steryl esters, causes the formation of lipid droplets. By electron microscopy, these lipid droplets are morphologically indistinguishable from those in wild-type cells. *In silico* and *in vitro* experiments confirmed the propensity of retinyl esters to segregate from membranes and to form lipid droplets. The hydrophobic N-terminus of LRAT displays preferential interactions with retinyl esters in membranes and promotes the formation of large retinyl ester-containing lipid droplets in mammalian cells. Our combined data indicate that the molecular design of LRAT is optimally suited to allow the formation of characteristic large lipid droplets in retinyl ester-storing cells.

## Introduction

Lipid droplets (LDs) form a ubiquitous class of organelles, best known for their role as storage of neutral lipids for a multitude of functions such as creating an energy reservoir, a source for building blocks, protection against lipotoxicity, a role in cell cycle, and storage of signaling lipids (Hashemi and Goodman, 2015; Pol et al., 2014; Thiam and Beller, 2017; Walther and Farese, 2012). However, LDs also play important roles in lipid metabolism and homeostasis. Dysfunction of LD synthesis has been linked to a range of diseases. The physiological role of LDs thus appears significantly larger than considered previously (Krahmer et al., 2013; Welte, 2015).

LDs have a unique organellar architecture with a lipid monolayer surrounding a hydrophobic core that consists of neutral lipids. A number of specific proteins associate with LDs regulating organelle and lipid dynamics. Recent advances have started to shed light on the mechanism of LD biogenesis. In the most prevalent view, LD formation is primarily driven by triacylglycerol (TAG) synthesis at the endoplasmic reticulum (ER). TAG accumulates at the interphase of the ER bilayer, until a critical demixing concentration is reached and phase separation occurs, leading to lens formation and membrane deformation (Thiam and Forêt, 2016; Walther et al., 2017). During this process of nucleation, neutral lipids coalesce to form lenses between the two leaflets of the membrane bilayer. Indeed, lenses of about 50nm have been observed in the ER upon induction of LD formation in yeast (Choudhary et al., 2015). As neutral lipid synthesis continues, nascent LDs may bud from the endoplasmic reticulum. Although this process may not require proteins other than TAG synthesizing enzymes such as DGAT 1/2 and ACSL3 (Kassan et al., 2013; Pol et al., 2014; Thiam and Forêt, 2016; Walther et al., 2017), several proteins and lipids have been identified in the regulation of LD number and size as well as LD dynamics.

The abundant presence of large LDs is a hallmark of hepatic stellate cells (HSCs) in normal liver. HSCs are specialized in the storage of retinol (vitamin A) as retinyl esters, giving the LDs their characteristic autofluorescent properties. After liver injury, the fine structure of HSCs changes considerably. They lose their characteristic LDs and transdifferentiate into myofibroblasts, in preparation to secrete collagen (Blaner et al., 2009; Friedman, 2008). Lipidomic analysis revealed complex dynamics of disappearance of different classes of neutral lipids during HSC activation (Testerink et al., 2012). Recent research shows the presence of two different types of LDs, so-called preexisting “original” LDs with relatively slow turnover and rapidly “recycling” LDs that transiently appear during activation of HSCs (Ajat et al., 2017; Molenaar et al., 2017; Tuohetahuntila et al., 2016). Whereas synthesis and breakdown of TAGs in rapidly recycling LDs is mediated by DGAT1 and ATGL (Tuohetahuntila et al., 2016), less is known about the turnover of preexisting LDs. Lysosomes play an important role in the degradation of these LDs (Tuohetahuntila et al., 2017) and this is likely to be related to the observed importance of the autophagic pathway in HSC activation (Hernandez-Gea and Friedman, 2011; Thoen et al., 2011).

Surprisingly, inhibition of DGAT1 does not affect the dynamics of the preexisting LDs nor does it affect the synthesis of retinyl esters in isolated primary HSCs (Ajat et al., 2017; Tuohetahuntila et al., 2016). However, HSCs contain a specialized enzyme called lecithin:retinol acyltransferase (LRAT) that catalyzes a trans-esterification reaction between the *sn-1* position of phosphatidylcholine (PC) and all-*trans*-retinol to form all-*trans*-retinyl ester (Fig. 1A,B) (Golczak et al., 2012; Ruiz and Bok, 2010). As LRAT is the main contributor to retinyl ester storage in the liver (Liu and Gudas, 2005; O’Byrne et al., 2005), we investigated the possibility that LRAT-mediated retinyl ester synthesis drives the generation of the relatively large, retinyl ester-containing LDs in quiescent HSCs.

**Figure 1.**
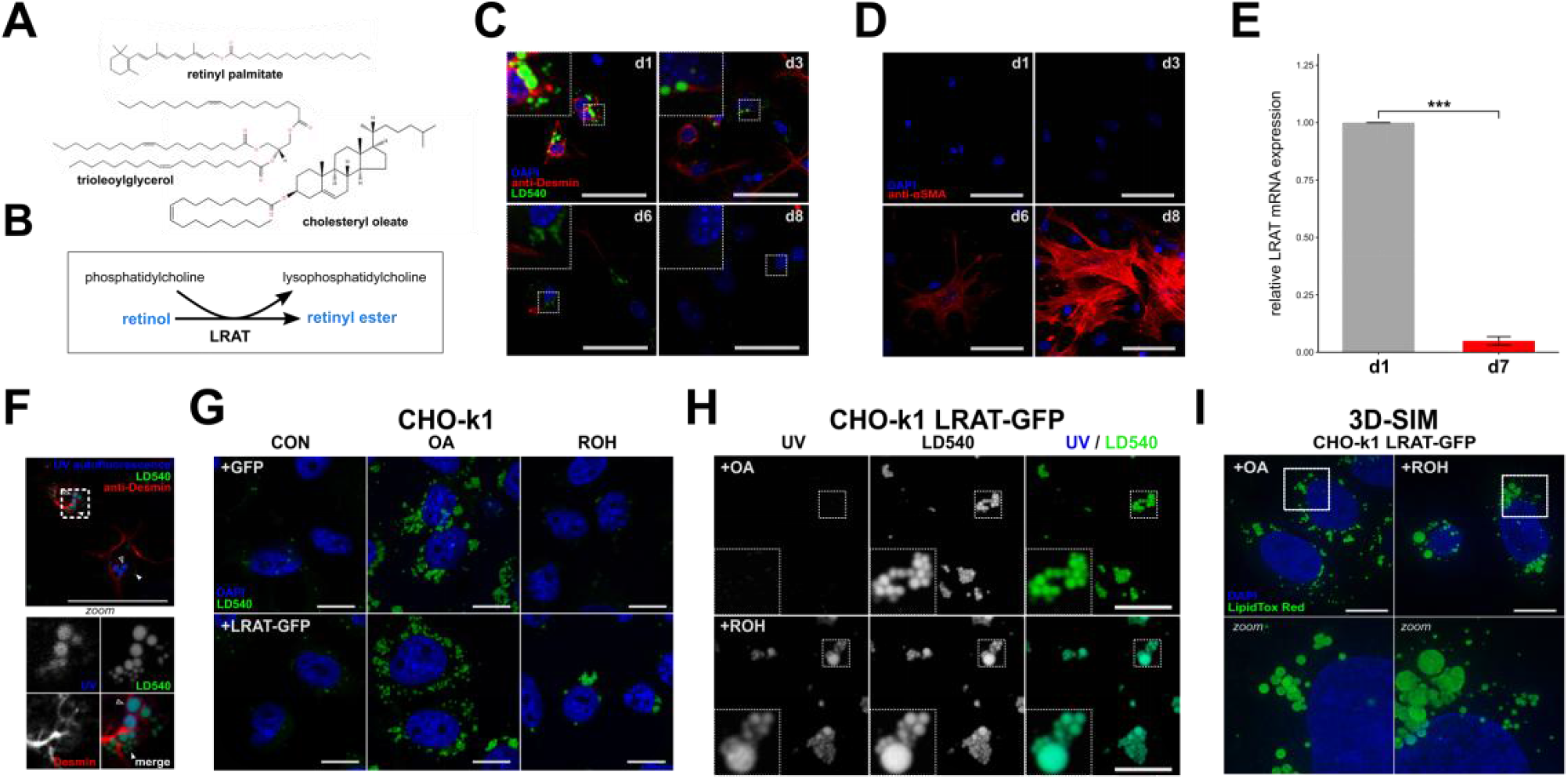
Lipid droplets containing retinyl esters have a distinct morphology. **(A)** Chemical structures of the neutral lipids retinyl palmitate, cholesteryl oleate, and trioleoylglycerol. **(B)** Reaction mechanism of LRAT. **(C,D)** Confocal microscopy of murine hepatic stellate cells (mHSCs) cultured for 1, 3, 6 or 8 days, stained with DAPI (blue), LD540 (green) and anti-desmin (red) **(C)**, or DAPI (blue) and anti-α-SMA (red) **(D)**. Scale bars indicate 50 μm. **(E)** Relative expression of LRAT mRNAin mHSCs after 1 or 7 days of culture by qPCR. Barplot indicates means ± SEM of 3 animals. Statistical significance was determined by a two-tailed paired Welch’s t-test. **(F)** Confocal microscopy of mHSCs two hours after isolation, showing UV-autofluorescence (blue), LD540 (green) and anti-desmin (red). Bottom panel is a zoomed inset of the area surrounded by dotted lines in the main panel. Closed triangles indicate UV^−^ LDs, open triangles indicate UV+ LDs. Scale bars indicate 50 μm. **(G,H)** Confocal microscopy of CHO-k1 cells expressing GFP or LRAT-GFP, incubated overnight in the presence or absence of 20 μM ROH or 200 μM OA. Imaged channels are DAPI (blue) and LD540 (green) **(G)**, or UV-autofluorescence (left) and LD540 (middle) **(H)**. Scale bars indicate 10μm. **(I)** Full projections of 3D-SIM images of CHO-k1 cells expressing LRAT-GFP, incubated overnight in the presence 20 μM ROH or 200 μM OA. Bottom panels are zoomed insets of areas surrounded by dotted lines in the top panels. Scale bars indicate 10μm. * P < 0.05, ** P < 0.01, *** P < 0.001, NS: not significant.

## Results

### LRAT expression generates UV-positive lipid droplets

Primary and quiescent HSCs spontaneously transdifferentiate into activated HSCs (myofibroblasts) *ex vivo* upon isolation and subsequent culture, resulting in LD disappearance. Quiescent and activated HSCs can be identified based on their high expression of desmin, whereas these two HSC populations can be distinguished from each other by an increased alpha smooth muscle actin (α-SMA) expression in activated HSCs (Blaner et al., 2009; Friedman, 2008) (Fig. 1C,D). In addition, LRAT expression decreases (Blaner et al., 2009; Kluwe et al., 2011) (Fig. 1E). We previously presented evidence for neutral lipid dynamics during HSC activation that is consistent with the existence of two different pools of LDs (Molenaar et al., 2017; Tuohetahuntila et al., 2017). To visualize these two different pools, we made use of the autofluorescent properties of retinyl esters in preexisting “original” LDs that are a hallmark of quiescent HSCs (Ajat et al., 2017; Friedman, 2008). After fixation of freshly isolated HSCs, retinyl esters (UV autofluorescence) and LDs (LD540) were imaged by confocal microscopy. We observed two distinct populations of LDs: large UV+LD540+ structures containing high amounts of REs and UV^−^LD540^+^ structures - depleted from REs - with smaller diameters (Fig. 1F). These observations are also in good agreement with our recent findings that LDs in LRAT^−/−^ HSCs were significantly smaller as compared to LDs from wild-type cells (Ajat et al., 2017). Together, these data suggest a role for LRAT in the formation of distinct vitamin A-containing LDs that are a hallmark of hepatic stellate cells (HSCs).

To understand the role of LRAT in LD biology, we stably transfected CHO-k1 cells with a plasmid carrying LRAT-GFP. Lipidomic analysis of CHO cells expressing LRAT showed that retinyl ester synthesis was observed only after addition of retinol (ROH) as substrate to the medium (Fig. 1G (UV^+^LDs) and Suppl. Fig. 1A) and revealed the presence of predominantly saturated (16:0 and 18:0) and mono-unsaturated (18:1) fatty acids (Suppl. Fig. 1A). This composition reflects the catalytic activity of LRAT, using fatty acids at the *sn-1* moiety of PC for esterification (Golczak et al., 2012; MacDonald and Ong, 1988). CHO cells do not contain detectable amounts of enzymatic activity of LRAT (Suppl. Fig. 1B) and in the absence of LRAT they can synthesize retinyl esters using a different mechanism involving DGAT1 (Ajat et al., 2017; Orland et al., 2005). This reaction occurs with much lower efficiency and results in a different species profile of retinyl esters (Suppl. Fig. 1C). Inhibition of DGAT1 activity in CHO-k1 cells expressing LRAT-GFP showed no inhibition of retinyl ester synthesis, which confirms that LRAT is the primary retinyl ester synthesizing enzyme (Suppl. Fig. 1D). Both CHO-k1 cells expressing GFP or LRAT-GFP showed increased LD540 fluorescence after incubation with oleic acid (OA). However, only LRAT-GFP expressing cells showed an increase in LDs after incubation with ROH, confirming the ability of LRAT to esterify ROH (Fig. 1G). Furthermore, these LDs exhibited UV-autofluorescence and were larger in diameter as compared to OA-stimulated LDs, which in turn were not autofluorescent (Fig. 1H). Similar results were obtained after transfection of human HSC-derived cell line LX-2 with LRAT (Suppl. Fig. 2). To exclude the possibility that the observed large UV^+^LDs were in fact clusters of small LDs that could not be distinguished due to the limited resolution of conventional confocal microscopy, we also imaged both conditions by super resolution microscopy (3D structured illumination microscopy, 3D-SIM). Using SIM, we observed both small and large sized LDs in LRAT-dependent LD-synthesis and with a clustered appearance (Fig. 1I). In the presence of OA, the LDs displayed a more homogeneous size-distribution of relative small LDs and a ‘dispersed’ localization through the cell (Fig. 1I, Suppl. Video 1-2).

### LRAT-mediated LD formation is independent of TAG synthesis

TAG synthesis is a driving force in LD biogenesis. The last step in TAG synthesis is performed by DGAT1 or DGAT2, enzymes that transfer activated fatty acids to diacylglycerol. Fatty acid activation is performed by acyl-CoA synthetases and their knockdown or pharmaceutical inhibition after OA-stimulation led to a decreased number of (pre-)LDs (Kassan et al., 2013). Acyl-CoA is, however, not involved in the transesterification reaction of a fatty acid from PC to ROH by LRAT to generate retinyl esters. To determine whether LRAT requires TAG synthesis for retinyl ester synthesis and formation of RE-containing LDs, we pre-incubated LRAT-GFP expressing cells under low serum conditions overnight and subsequently incubated the cells in the presence of triacsin C, a drug that inhibits acyl-CoA synthesizing activities (Igal et al., 1997). In the presence of triacsin C, the number of LDs was strongly reduced, both in the presence and absence of OA (Fig. 2A). In the presence of ROH, however, formation of large LDs was still observed. These results confirm that LRAT does not depend on the presence of acyl-CoA and supports the possibility that LD-formation via LRAT has a distinct mechanism that is independent of TAG (DGAT/ACSL-mediated) LD formation.

**Figure 2.**
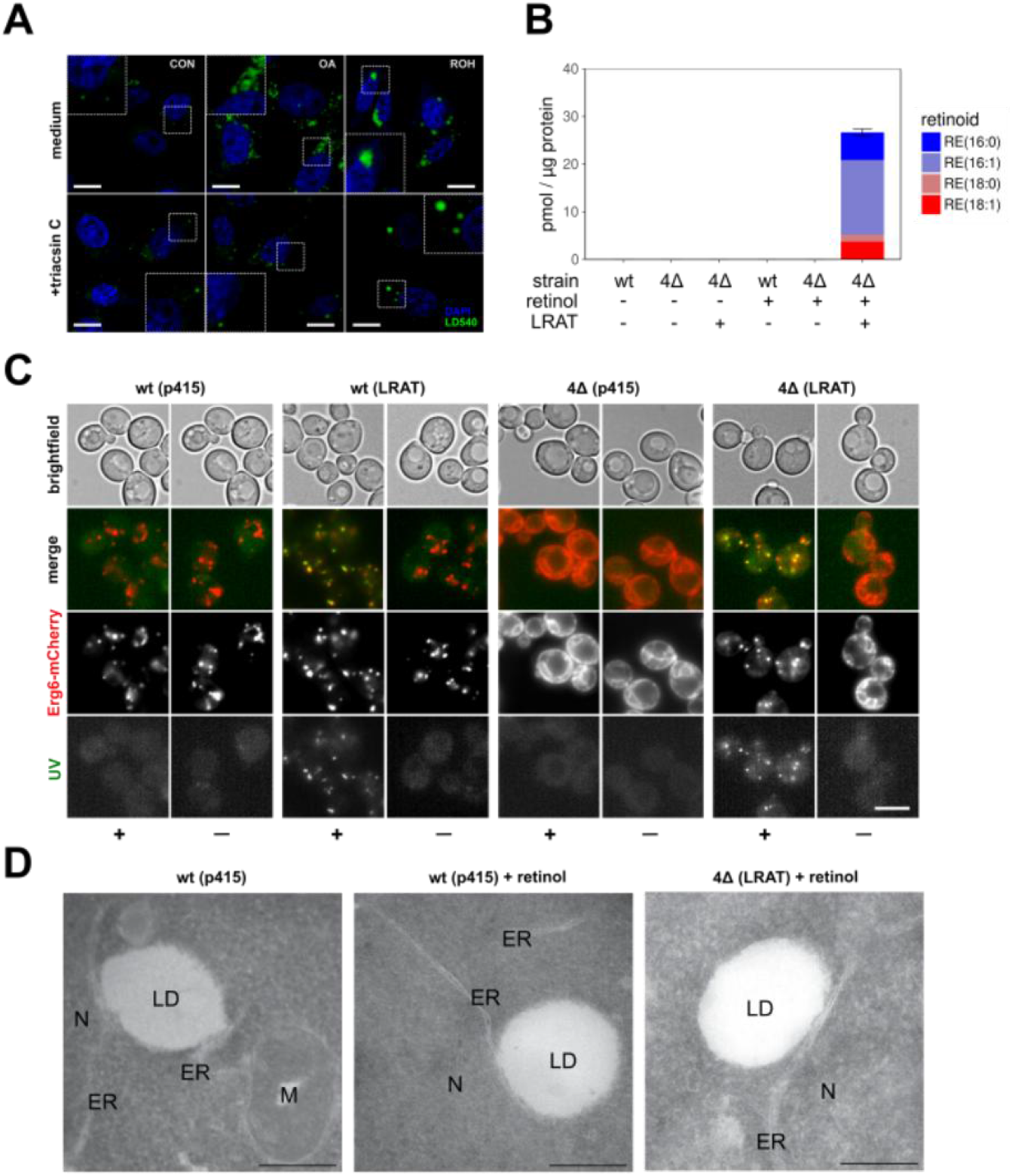
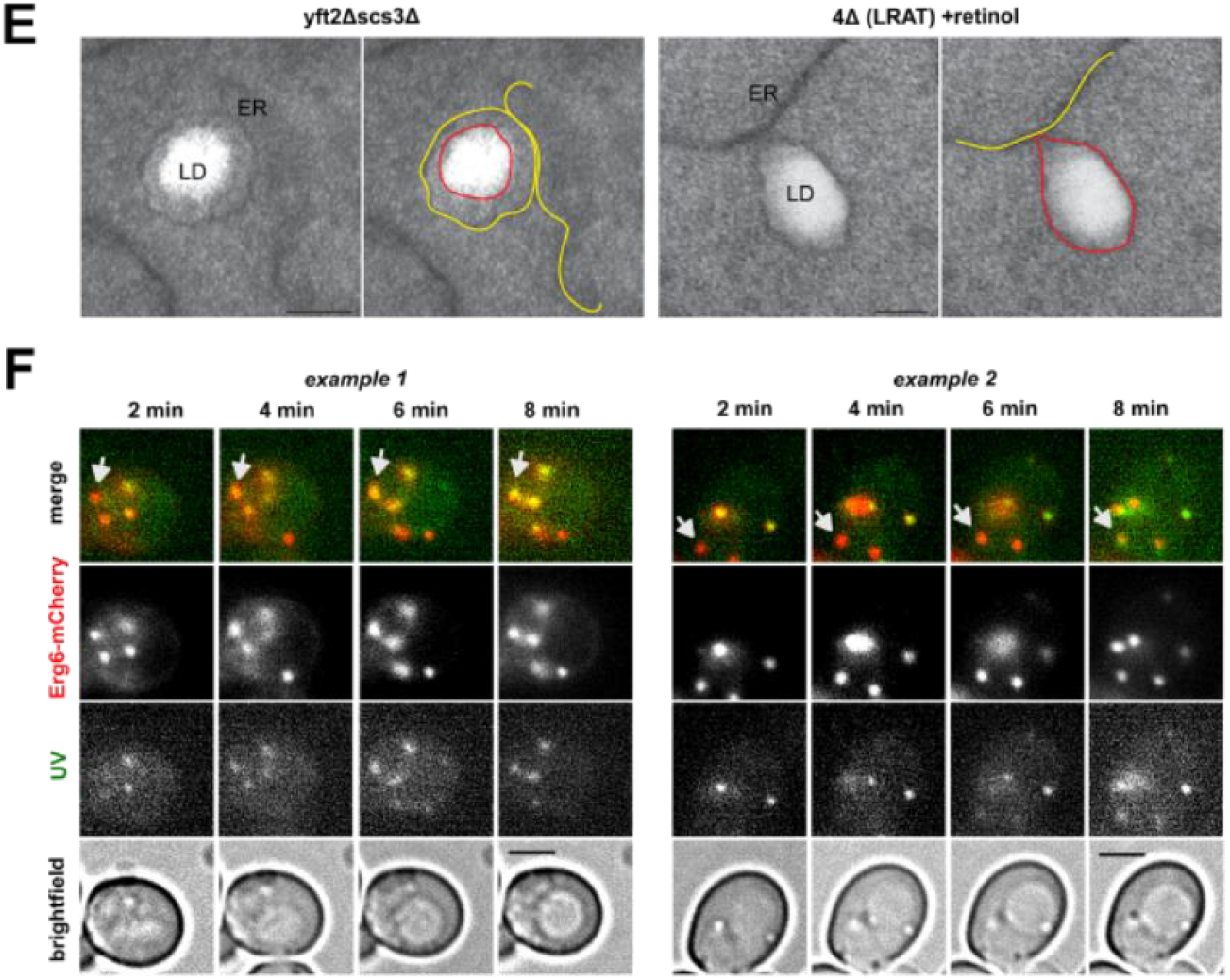
LRAT-mediated LD-formation in the absence of pre-existing LDs. **(A)** Confocal microscopy of CHO-k1 expressing LRAT-GFP. After pre-incubation with medium containing 1% FBS, cells were incubated overnight with or without 200 μM OA or 20 μM ROH, in the presence or absence of 1 μg/mL triacsin C. Cells were stained with DAPI (blue) and LD540 (green). Scale bars indicate 10 μm. **(B)** Quantification of predominant RE-species by LC-MS/MS of wt and 4Δ yeast cells expressing LRAT, 2 hours after incubation in the presence or absence of 2 mM ROH. Amounts in pmol RE per μg protein. Barplot indicates means ± SD. **(C)** Wide-field microscopy of Erg6-mCherry expressing wt and 4Δ yeast cells, with or without expressing LRAT, 2 hours after incubation with or without 2 mM ROH (‘—’ or ‘+’). Images of UV-autofluorescence (green), Erg6-mCherry (red) and brightfield were taken. Scale bars indicate 5 μm. **(D)** Electron microscopy of wt and 4Δ yeast cells, with or without expressing LRAT, incubated with or without 2 mM ROH. ER, endoplasmic reticulum; LD, lipid droplet; M, mitochondria; N, nucleus. Scale bars indicate 250 nm. **(E)** Electron microscopy of yft2Δscs3Δ, and 4Δ yeast cells expressing LRAT, incubated with 2 mM ROH. Scale bars indicate 100 nm. ER, endoplasmic reticulum (yellow lines in right panels); LD, lipid droplet (red lines right panels). **(F)** Wide-field microscopy of Erg6-mCherry expressing wt yeast cells expressing LRAT. Two time series of UV-autofluorescence (white), Erg6-mCherry (red) and brightfield, taken 2, 4, 6 and 8 minutes after addition of 2 mM ROH. Scale bars indicate 5 μm.

Some LDs could still be observed in cells treated with triacsin C and therefore we could not exclude the possibility that, rather than TAG synthesis, TAG-filled LDs are required for RE-containing LD formation. Therefore, we made use of another model system, a *Saccharomyces cerevisiae* mutant strain that lacks the four enzymes responsible for the last steps in triglyceride - Lro1 and Dga1 - and steryl ester (SE) - Are1 and Are2 - synthesis. This mutant strain is viable, but does not contain LDs (Sandager et al., 2002). After introduction of human LRAT into *IrolΔ dgalΔ arelΔ are2Δ* cells (hereafter 4Δ cells), LRAT-GFP co-localizes with Sec63-mCherry, an ER marker in yeast (Suppl. Fig. 3A). The resulting LRAT expressing yeast cells (4ΔLRAT) were able to synthesize REs after the addition of ROH (Fig. 2B). In contrast to mammalian cells, the predominant RE-species was retinyl palmitoleate, RE(16:1). This is in line with the reported fatty acid composition of PC in yeast, which is capable of producing only monounsaturated fatty acids, and contains predominantly PC(32:2) and PC(34:2) (Boumann et al., 2006). In the absence of either LRAT or ROH, both 4Δ and wild-type (wt) yeast cells did not synthesize REs. These results also demonstrate the absence of an endogenous RE synthase activity in yeast by acyl-CoA retinol acyltransferase (ARAT) activity e.g. Dga1 (Fig. 2B). Thus, RE synthesis in yeast depends on LRAT expression. As anticipated, 4Δ LRAT yeast cells were devoid of LD structures in the absence of exogenous ROH, as evidenced by membrane localization of LD-marker Erg6 (Fig. 2C) and BODIPY (Suppl. Fig. 3B). Upon addition of ROH to 4ΔLRAT yeast cells, we observed a clear presence of autofluorescent LD-like structures and co-localization of these structures with the LD-marker Erg6 (Fig. 2C, Suppl. Fig. 3C) and with BODIPY (Suppl. Fig. 3B). In the absence of LRAT, 4Δ yeast cells could not generate LDs in the presence of ROH.

Electron microscopic examination of the LDs generated in 4Δ LRAT yeast cells revealed the presence of *bona fide* LDs with a cytosolic orientation that were morphologically indistinguishable from TAG/SE-filled LDs generated in wt yeast cells (Fig. 2D,E). These results demonstrate that LRAT induces the formation of LDs in the absence of other LDs or TAGs (Suppl. Fig. 4). To determine whether retinyl esters can partition into existing LDs or exclusively form their own LDs, we determined the co-localization of RE positive LDs (UV^+^) with all LDs (Erg6^+^) immediately after addition of ROH to *wt* yeast cells expressing LRAT. These cells have TAG-filled LDs present before addition of ROH. As shown in Fig. 2F, 2 min after addition of ROH, co-localization of Erg6^+^ LDs with UV^+^ LDs was observed. In addition, UV^−^ LDs become UV^+^ in the next 6 min, indicating that REs are also transferred into existing LDs.

### Spontaneous nucleation and lens formation of retinyl esters in lipid bilayers

The mechanism of LD formation by LRAT-mediated RE synthesis is not known. For TAG-filled LDs it has been shown that TAGs have a limited solubility in biological membranes. Above a critical demixing concentration (∼3%), TAG molecules segregate to form TAG lenses that appear as a first step in LD formation (Choudhary et al. 2015; Khandelia et al., 2010; Thiam and Forêt, 2016). After this ‘nucleation’ process, the lenses grow and can emerge from ER membranes during a ‘budding’ process. The formation of lenses in TAG containing membranes has been demonstrated and studied in molecular dynamics (MD) simulations (Ben M’barek et al., 2017; Khandelia et al., 2010). Here we built upon that approach and used coarse-grain MD (CG-MD) simulations of 1-palmitoyl-2-oleoyl-sn-glycero-3-phosphocholine (POPC) membranes with different amounts of trioleoylglycerol (TOG) and retinyl palmitate (RP) to assess the propensity of lens formation in those systems. In setups containing only TOG, lens-formation was consistently observed and always completed within 100 ns of simulation. In contrast, setups with only RP took considerable longer to nucleate (Fig. 3A) and more often failed to form lenses on the time scale used for the simulations (250 ns). Lenses formed by RP were also typically less well defined, with more RP remaining dispersed throughout the membrane. Mixtures of TOG and RP showed an intermediate efficiency of lens-formation (Fig. 3A and B). These simulations imply that RP has a lower propensity to self-aggregate, and hence has a higher nucleation barrier as compared to TOG.

**Figure 3.**
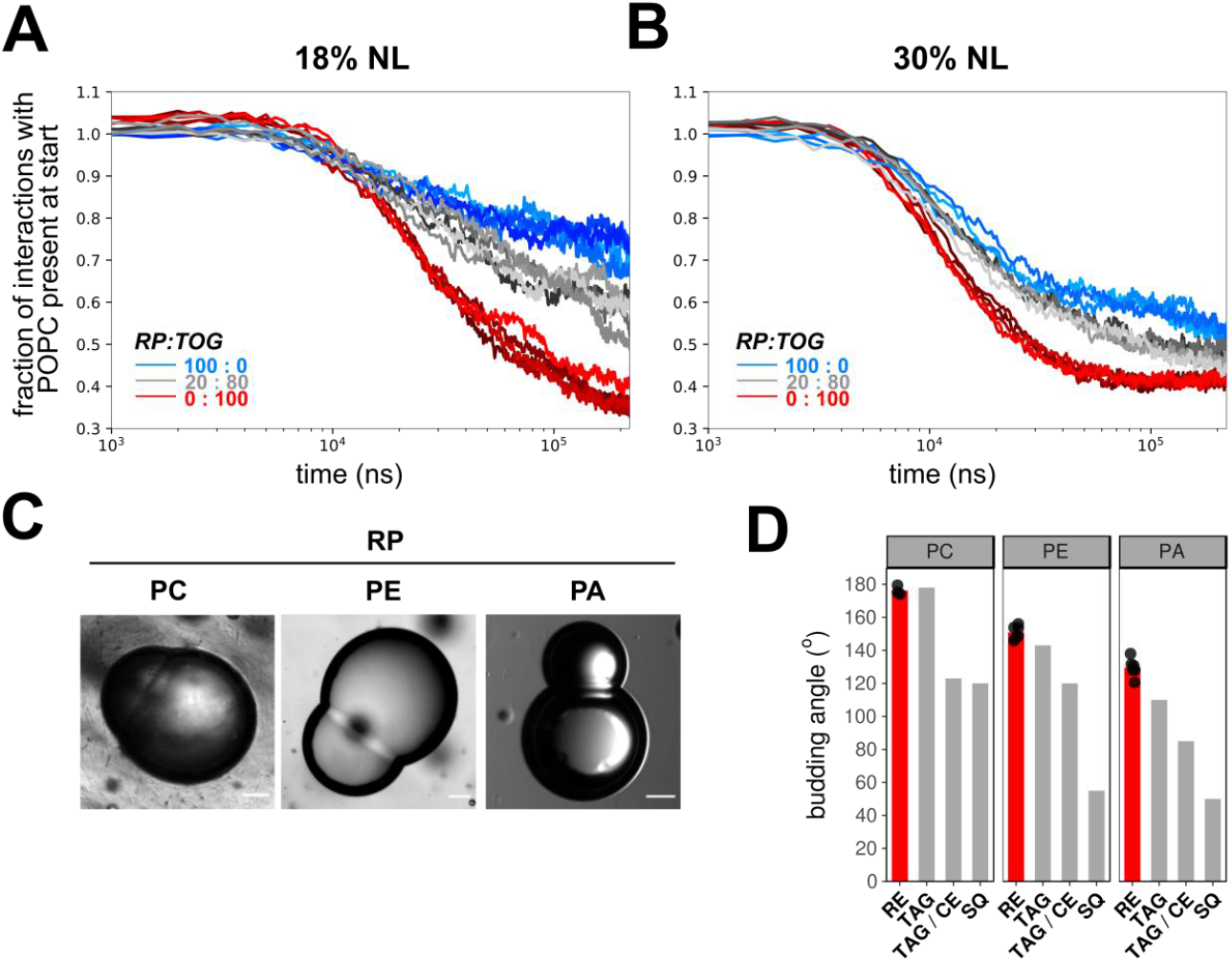
Nucleation and budding properties of RE. **(A,B)** Coarse grain MD simulations of lens formation by 150 (18%) **(A)** and 250 (30%) **(B)** neutral lipids per leaflet in a POPC membrane. Colors indicate the neutral lipid composition, with marine for pure RE and red for pure TOG. The progress of lens formation is shown as fractional loss of interactions between the neutral lipids and POPC as function of time (log-scale). **(C)** Brightfield microscopy images of droplet interface bilayers of droplets containing neutral lipid RP and lipid surfactants PC, PE or PA. Scale bars indicate 20 μm. **(D)** Comparison of quantified budding angles (mean±SD) of RP-containing droplets with lipid surfactants PC (176,2±2,8), PE (150,9±4,0) or PA (129,5±5,7) vs. reported budding angles of TAG and SQ-containing droplets (Ben M’barek et al., 2017). Barplot represent means of at least 8 individual measurements, which are shown as black dots.

Methods to directly measure the nucleation barrier (or ‘critical nucleation volume’) do not exist, but in a setting with the same PL-composition, the monolayer surface tension will be a driving force that affects nucleation (Ben M’barek et al., 2017; Deslandes et al. 2017; Thiam and Forêt, 2016). The deformation and budding of a LD monolayer is controlled in part by the monolayer bending rigidity and surface tension – the monolayer bending rigidity will tend to flatten a lens, thereby re-dispersing neutral lipid molecules in the bilayer, while its surface tension will tend to make a lens spherical. However, at a characteristic size, surface tension will essentially control the monolayer deformation. This size will be larger for a lower monolayer surface tension: lower monolayer tensions will likely result in larger ‘critical nucleation volumes’ or higher nucleation barriers (Ben M’barek et al., 2017; Chorlay and Thiam, 2018; Deslandes et al., 2017; Thiam and Forêt, 2016). We measured the tension of artificial LD containing either RP or TOG and, in line with our MD simulations, tensions were considerably lower in droplets made of RP as compared to their TOG counterparts (Table 1). Moreover, droplets surrounded by DOPC with maximum phospholipid packing showed the same trend. These measurements suggest that RP droplets have higher nucleation barriers, in agreement with the results obtained by CG-MD.

**Table 1.** Interfacial tension values by drop tensiometry

| system | tension (mN/m) |
| --- | --- |
| TOG / H<sub>2</sub>O | 32 |
| RP<sup>1</sup> / H<sub>2</sub>O | 8-14 |
| TOG / DOPC / H<sub>2</sub>O | 0.5-0.6 |
| RP<sup>1</sup> / DOPC / H<sub>2</sub>O | 0.08-0.2 |

<sup>1</sup>based on density provided by manufacturer, 0.90-0.95 g/mL

The efficiency of the subsequent budding process of RE-containing LDs can be studied by determination of budding angles in droplet-embedded vesicles or droplet interface bilayers (DIBs) containing neutral lipids (Ben M’barek et al., 2017; Chorlay and Thiam, 2018). By using the DIB system, we compared the budding angles of RP-containing lipid phase with reported values of TAG-LDs (Fig. 3C,D). The budding angles of RP^+^ droplets with PC and PE monolayers did not differ from reported values of their TOG^+^ counterparts (Fig. 3D); angles in the case of PA were only slightly higher. These data suggest that RP forms spontaneously LDs as efficiently as TG. In PC tensionless membranes, smaller RP-LD diameters would be thus expected, based on the budding angles, but the delay in nucleation might introduce an increase in the droplet size. This is, however, not observed in yeast cells (Fig. 2C). Thus, other factors must contribute to the characteristic large size of retinyl ester-containing LDs.

### N-terminal sequence of LRAT affects lipid droplet morphology

To study a role of LRAT in LD morphology, we considered the possibility that the N-terminus of LRAT aids in the formation of LDs. The N-terminal domain of LRAT (hereafter, LRAT-Nt AH) does not form a trans-membrane domain, despite its high hydrophobicity (Moise et al., 2007). In addition, it was reported that this domain can localize to LDs (Jiang and Napoli, 2012). We stably transfected CHO-k1 cells with a GFP fusion construct of LRAT lacking the N-terminus (ΔNt-LRAT-GFP) and selected clones with similar GFP fluorescence as compared to CHO-k1 cells stably transfected with full-length LRAT-GFP (Fig. 1). LRAT-activity of both homogenates was analyzed *in vitro* using HPLC-MS/MS by determination of RE(7:0) synthesis upon addition of exogenous PC(7:0/7:0) (Fig. 4A, see Materials and Methods for details). As expected, both enzymes displayed LRAT-activity with similar Km-values for ROH, but reaction rates of ΔNt-LRAT-GFP homogenates were somewhat lower (Fig. 4B). After overnight incubation with ROH, we observed UV^+^ LDs in both cell lines, but LDs appeared smaller in size in ΔNt-LRAT-GFP expressing cells (Fig. 4C,D). These data show that the N-terminus is not required *per se* for LD generation, but may have an effect on LD morphology. To rule out the possibility that the reduced LD size in the presence of ΔNt-LRAT-GFP is caused by the reduced activity of this LRAT mutant (Fig. 4B), we developed a single-cell imaging analysis pipeline. To this end, we determined cell size, LD-number per cell and mean LD size per cell by image analysis (see Materials & Methods for details). Total LD volume (representative for total neutral lipid content) was calculated from these parameters. Using this approach, we were able to select (“gate”) cells with similar total LD volume and thus neutral lipid content per cell (Fig. 4E). We then compared the log-ratios of LD number *vs*. LD volume per cell of both cell lines incubated with OA or ROH (Fig. 4F). As expected, no difference was observed when both cell lines were incubated with OA (Fig. 4F, left panel). In contrast, LDs generated in the presence of ROH were smaller in size and larger in number in cells expressing ΔNt-LRAT-GFP as compared to LDs from cells expressing the full-length protein, as reflected by decreased log-ratios (Fig. 4F, right panel). This difference in LD size:number distribution could also be observed in ungated cells; expressing LD-number as function of the total LD volume per cell showed that regression of all datapoints of LRAT lacking its N-terminus results in a steeper slope as compared to full-length LRAT, indicating more LDs per amount of total neutral lipid (Fig. 4G). Taken together, these results show that the LRAT-Nt has the ability to affect the size:number distribution of retinyl ester-containing LDs in mammalian cells.

**Figure 4.**
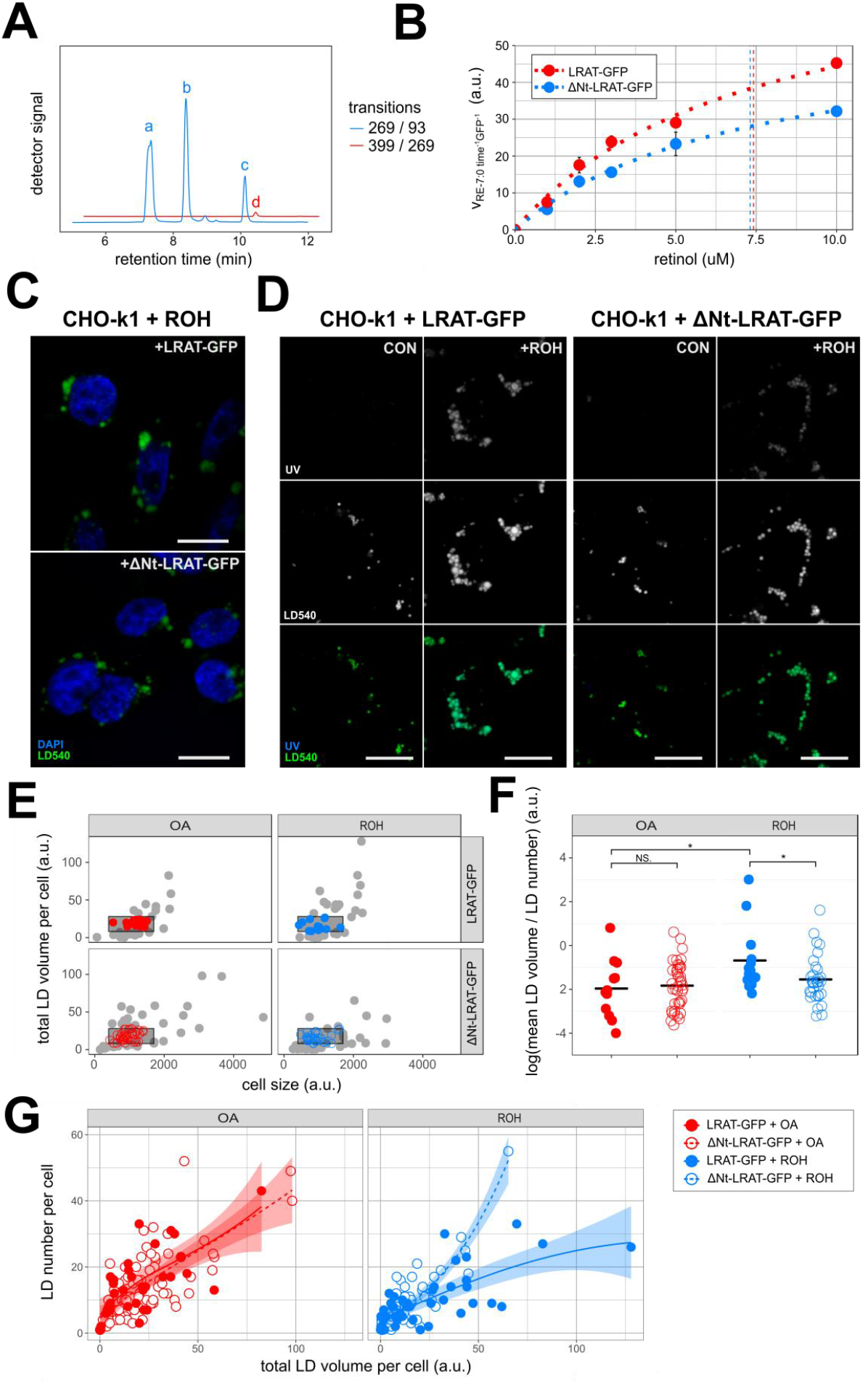
Deletion of N-terminus of LRAT affects LD formation in CHO cells. **(A,B)** *In vitro* LRAT-activity of CHO-k1 homogenates expressing full-length or ΔNt-LRAT-GFP. After incubation full-length LRAT-GFP with PC(7:0/7:0) and varying concentration of ROH, ROH (peak a), RE(2:0) (internal standard, peak b) and RE(7:0) (peak c) were measured by LC-MS/MS. MRM-transition 269/93 detects all retinoid backbones (blue), whereas 399/269 specifically detects RE(7:0) (red) **(A)**. RE(7:0)-synthesis by full-length or ΔNt-LRAT-GFP was normalized by GFP-levels of the homogenates and plotted against increasing ROH concentrations. Estimated Km values are indicated by vertical dashed lines. A representative plot of three independent experiment is shown **(B)**. **(C,D)** Confocal microscopy of CHO-k1 cells expressing LRAT-GFP or ΔNt-LRAT-GFP, incubated overnight in the presence or absence of 20 μM ROH. Imaged channels are DAPI (blue) and LD540 (green) **(C)**, or UV-autofluorescence (top) and LD540 (middle) **(D)**. Scale bars indicate 10 μm. **(E,F)** Quantification of cells imaged by confocal microscopy. Cell sizes were plotted against total LD volume per cell (LRAT-GFP or ΔNt-LRAT-GFP) and cells with similar total LD volumes per cell were gated (black box, LRAT-GFP+OA: 12 cells; ΔNt-LRAT-GFP+OA: 40 cells; LRAT-GFP+ROH: 13 cells; ΔNt-LRAT-GFP+ROH: 31 cells) **(E)**. The gated cells were expressed as log-ratios of mean LD volume vs. LD number. Mean values (±SD) were −1,96±1,34 (LRAT-GFP with OA); −1,83±1,06 (ΔNt-LRAT-GFP with OA); −0,68±1,51 (LRAT-GFP with ROH); and −1,55±1,08 (ΔNt-LRAT-GFP with ROH) Statistical significance was determined by two-tailed unpaired Student’s t-tests, P-values were corrected by the Benjamini-Hochberg procedure **(F)**. Cells were incubated overnight with 200 μM OA (red circles) or 20 μM ROH (blue circles) and LD number per cell as a function of total LD volume per cell for LRAT-GFP (closed circles) or ΔNt-LRAT-GFP-expressing (open circles) was analyzed without gating **(G)**. Every data point represents one cell. Lines are moving averages, shades indicate 95% confidence intervals. LRAT-GFP+OA: 35 cells; ΔNt-LRAT-GFP+OA: 79 cells; LRAT-GFP+ROH: 43 cells; ΔNt-LRAT-GFP+ROH: 68 cells. * P < 0.05, ** P < 0.01, *** P < 0.001, NS: not significant.

### N-terminus of LRAT exhibits affinity for retinyl esters *in silico*

The N-terminus of LRAT is predicted to have an alpha-helical structure without hairpin topology, which is flanked by amino acids without a clear secondary structure (Suppl. Fig. 5). HeliQuest analysis (Gautier et al., 2008) showed a clear separation between hydrophobic and polar residues on both sides of the helix, in line with an amphipathic topology (Fig. 5A, upper panel). However, in comparison with canonical amphipathic helices involved in membrane recognition (ALPS motifs) (Bigay et al., 2005), the fraction of hydrophobic over polar residues was considerably higher in the LRAT N-terminal alpha-helix. This is exemplified by comparison of this projection with CCT-α P2, a recently characterized amphipathic domain with LD-affinity (Prévost et al., 2018) that contains an ALPS-like architecture (Fig. 5A, bottom panel). The helical wheel projection of the CCT-α P2 AH revealed a smaller fraction of hydrophobic residues as compared to the LRAT-Nt AH. Membrane-binding of CCT-α P2 is proportional to the amount of lipid packing defects (Prévost et al., 2018). We compared the membrane-binding characteristics of LRAT-Nt with CCT-α P2 by CG-MD using the DAFT-approach (ensemble of 30 simulations per combination of peptide and model membrane) (Wassenaar et al., 2015a). One model membrane consisted of POPC and the other one contained POPC and dioleylglycerol (DOG), a lipid that induces packing defects (Vamparys et al., 2013). We subsequently compared membrane binding of both LRAT-Nt and CCT-α P2. The binding half-time of CCT-α P2 to POPC membranes was 150 ns (Fig. 5B, bottom panel) and in the presence of DOG, this binding was accelerated (50 nsec) (Fig. 5B, bottom panel), confirming the role of packing defects in membrane binding (Prévost et al., 2018). For LRAT-Nt, however, we did not observe a difference in the kinetics of membrane binding between the two model systems (Fig. 5B, top panel; median of about 60 nsec). Interestingly, the results also showed that LRAT-Nt AH docked deeper into the bilayer than the CCT-α P2 AH (Fig. 5C), suggesting that LRAT-Nt has the potential to directly interact with neutral lipids, whereas CCT-α P2 AH binds only to phospholipids.

**Figure 5.**
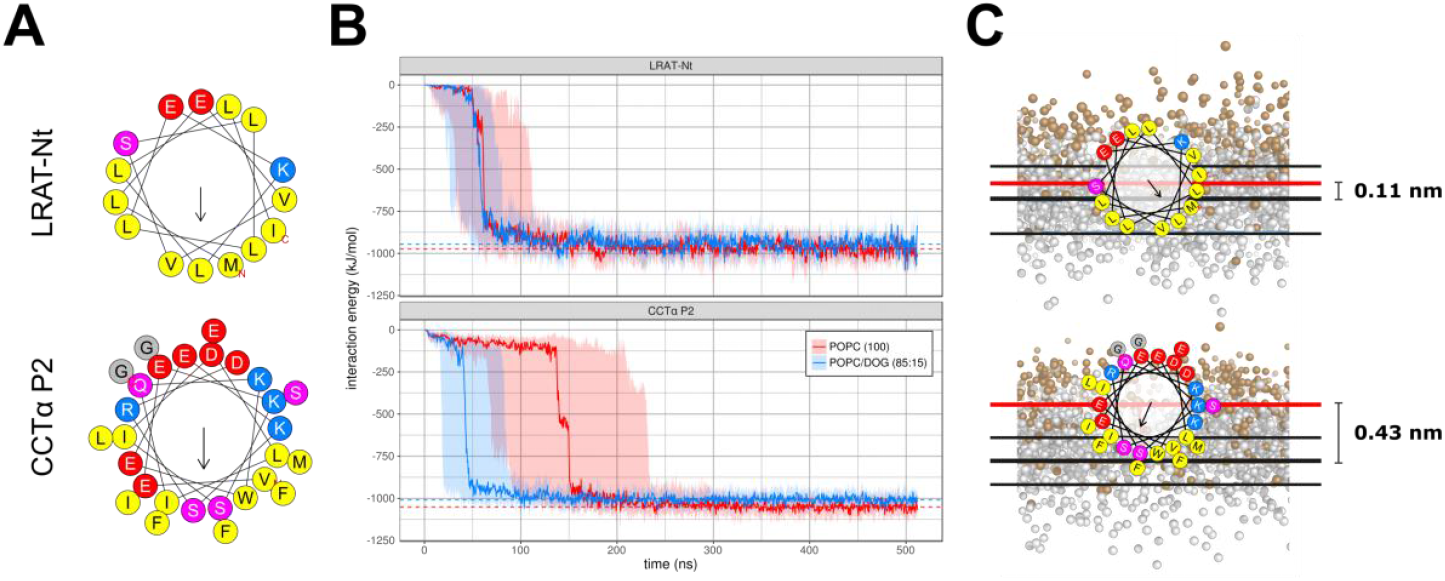
Amphipathic properties of LRAT-Nt and CCT-α P2 helices. **(A)** Helical wheel projections by HeliQuest of LRAT-Nt (top) and CCT-α P2 (bottom). Colors indicate amino acid categories: hydrophobic (yellow), negatively charged (red), polar (purple), positively charged (blue) and other (gray) residues. The hydrophobic moment is indicated with a black arrow. **(B)** Peptide-membrane interaction energies for LRAT-Nt (top) and CCT-α P2 with pure POPC (red) and with an 85:15 POPC/DOG mixture (red) over time across DAFT simulation ensembles of 30 simulations each, showing the region from the 1st to 3rd quartile as shaded areas and the median interaction energy for each ensemble of simulations as solid line. **(C)** Insertion depth of LRAT-Nt and CCT-α P2, showing the helical wheels of LRAT-Nt (top) and CCT-α P2 according to their mean angular orientations, with a red line signifying the position of the helix center, the thick black line denoting the median position of the lipid head groups across simulations and the thinner line denoting the standard deviation of the median across simulations. Colors in the peptides are as described in panel A.

To investigate a potential interaction of the N-terminal peptide with neutral lipids, we extended the MD setup to a series of POPC membranes containing 10% neutral lipids in different ratios RP:TOG, and in the presence of LRAT-Nt. The simulations showed that LRAT-Nt typically co-localized with neutral lipids (Fig. 6A). Inspection of the localization of LRAT-Nt with respect to the membrane profile (Fig. 6B; Suppl. Fig. 6A) confirmed that the peptide is docked deep into the membrane. Surprisingly, LRAT-Nt is docked deeper into the membrane when the neutral lipids consist predominantly of RP as opposed to TOG (Fig. 6B). This is only partly explained by the thicker membrane resulting from the more complete lens formation at higher TOG ratios. Initial inspection of the localization of TOGs and RPs within the lens revealed an asymmetric distribution of the neutral lipid mixture with RPs in closer proximity to the LRAT-Nt peptide (Suppl. Fig. 6B). Assessment of the interaction energies between the LRAT-Nt helix and both RP and TOG confirmed that the helix has stronger interactions with RP than it has with TOG (Fig. 6C). This specific affinity of LRAT-Nt for REs could affect lens and/or LD formation by REs, for example by facilitating the formation of RP containing lenses by decreasing the nucleation energy barrier. This should then be reflected in an enhanced nucleation rate. To study such an influence on lens formation, we performed several series of MD simulations. However, under none of these conditions LRAT-Nt significantly enhanced the rate of lens formation (data not shown).

**Figure 6.**
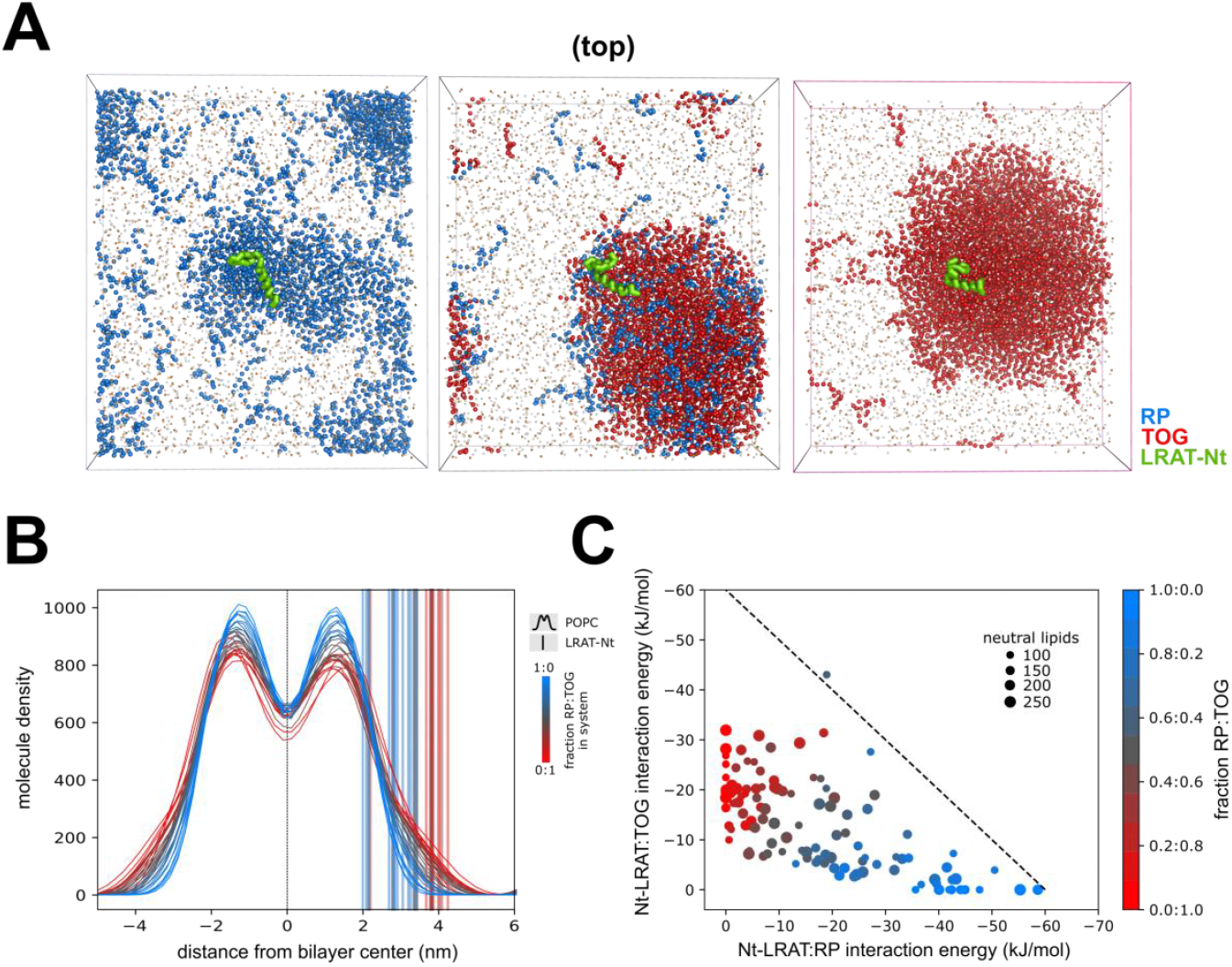
Affinity of LRAT-Nt for retinyl esters in GC-MD. **(A)** Binding of LRAT-Nt to lenses consisting of 300 molecules RP (left), 120+180 molecules RP+TOG (middle) and 300 molecules TOG (right). The panels show preferential binding of the helix (green) to the region of maximal curvature. The RP and TOG particles are shown in marine and red, respectively. **(B)** Positions of centers of mass of LRAT-Nt with respect to the membrane as function of the ratio RP:TOG (marine: pure RP to red: pure TOG). The membrane is shown as density profile, while the positions of LRAT-Nt are marked with dashed lines. **(C)** Specificity of interaction of LRAT-Nt with RP over TOG. Each simulation represents a dot and indicates the interaction energy between LRAT-Nt and RP on the x-axis and between LRAT-Nt and TOG on the y-axis. Dot size reflects the total number of neutral lipids and the color signifies the ratio between RP and TOG as indicated. The dashed line has slope −1 and a parallel profile is indicative of a transition from one pure compound to the other with no preferential interactions. The observations show a lower interaction with TOG as compared to RP, resulting in a profile with a reduced slope.

### N-terminus of LRAT exhibits affinity for retinyl esters *in vitro*

To study the affinity of the LRAT N-terminus to RP *in vitro*, we synthesized the LRAT-Nt peptide and assessed its recruitment to RP vs. TOG interface. Oil-in-water droplets were formed and the peptide was added (Fig. 7A). Recruitment of the peptide to the droplet interface will decrease the interfacial surface tension (Fig. 7A) (Ajjaji et al., 2019; Small, Wang, & Mitsche, 2009), which happened for both RP and TOG (Fig. 7B). Tension reached an equilibrium value which corresponds to the maximum of adsorbed peptides (9.2 mN/m for TOG, and 6.1 mN/m for RP). The peptide recruitment level is determined by the neutral lipid affinity for LRAT-Nt. To further investigate this, we performed a rapid compression experiment whereby, at the equilibrium interfacial tension, the droplet interface was reduced by decreasing its volume (Fig. 7C,D, left, dotted lines). This operation transiently increased the interfacial lateral pressure, *i.e*. the peptide surface density, and we recorded the relaxation of the system to equilibrium. During compression, surface tension decreased in the case of TOG, reached a transient plateau and subsequently continued decreasing (Fig. 7C). Appearance of the plateau is a signature of a rearrangement of the peptide at the surface, induced by the increase of its surface density (Mitsche & Small, 2013). As soon as compression was stopped, surface tension increased again, which is a signature of the desorption of some of the peptides from the surface. This behavior at the TOG interface is unique to LRAT-Nt, as the 11mer repeat domain of Plin1 does not show such response under similar experimental procedures (Ajjaji et al., 2019). With RP, compression led to a continuous decrease of tension (Fig. 7D) and upon arrest of compression, tension remained constant: the peptide did not desorb from the interface. These results confirm the MD simulations and show that the LRAT-Nt peptide interacts with neutral lipids and with a preference for retinyl esters.

**Figure 7.**
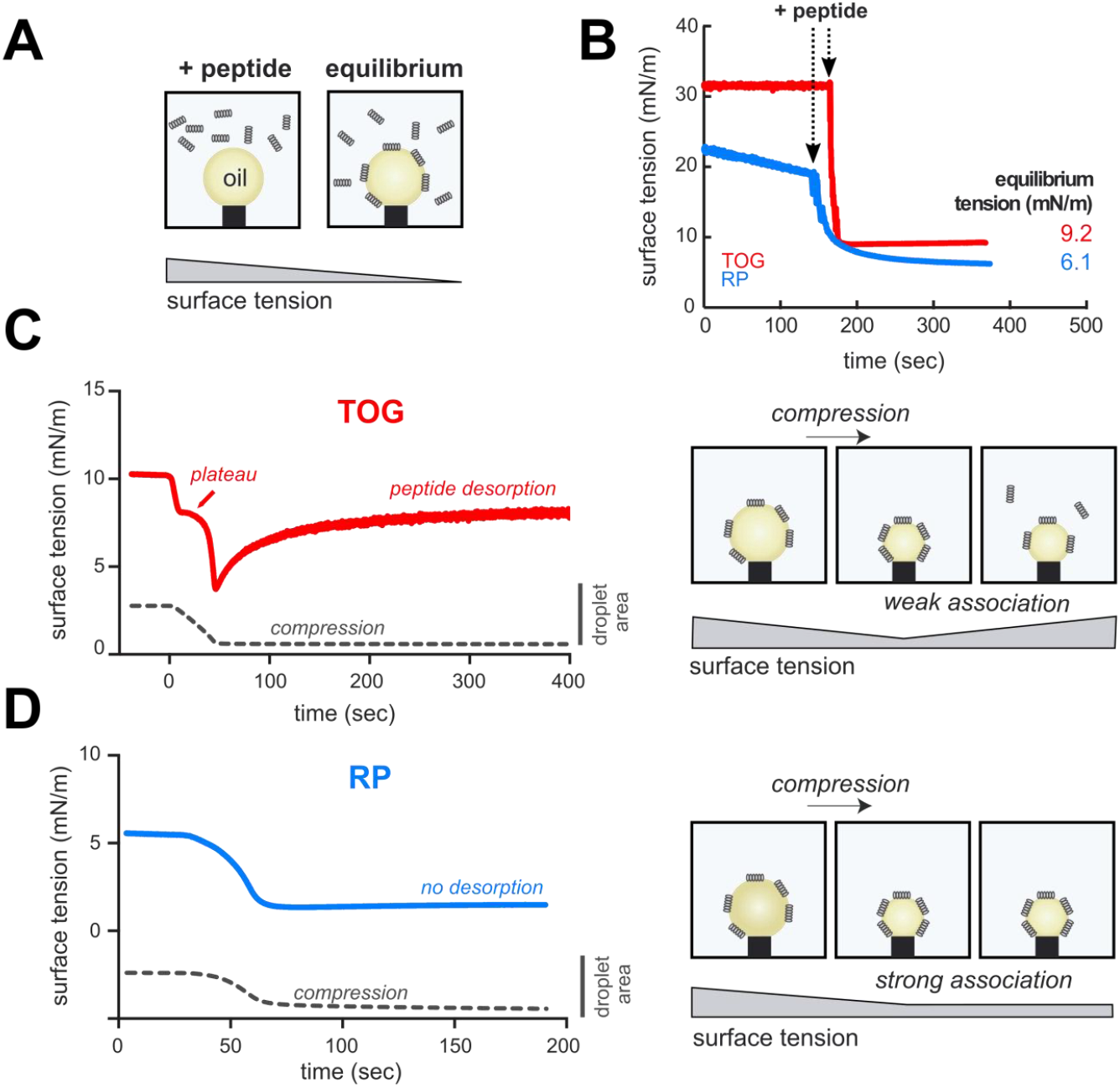
LRAT N-terminal peptide shows strong association with retinyl palmitate droplets. **(A)** Left, peptide addition to the buffer surrounding a pending oil droplet, trioleoylglycerol (TOG) or retinyl palmitate (RP), results in a decrease of the interface surface tension induced by peptide recruitment. **(B)** Related surface tension quantification over time for TOG and RP oil droplets. Arrows indicate time of LRAT-Nt peptide injection in the buffer. **(C)** After, equilibrium is reached, the surface of the TOG-droplet covered by the peptide is compressed, provoking a drop of surface tension. When compression was stopped, surface tension re-increased, indicating desorption of the LRAT-Nt peptide from the TOG-buffer interface. Right panel illustrates the manipulations and consequence on tension and peptide behavior. **(D)** After the LRAT-Nt peptide was recruited to the RP-droplet surface, the interface was compressed, provoking a surface tension drop. When compression was stopped, surface tension remained constant, indicating a strong association of the peptide with the RP-buffer interface. Right panel illustrates the manipulations and consequence on tension and peptide behavior.

## Discussion

Lipid droplet formation starts when neutral lipids accumulate within the bilayer of biological membranes above the a critical demixing concentration (Thiam and Forêt, 2016; Walther et al., 2017). Although the intrinsic capacity of different classes of neutral lipids to form LDs is well accepted based on biophysical considerations (Ben M’barek et al., 2017), *in vivo* this has only been demonstrated for triacylglycerols in yeast (Sandager et al., 2002). As cholesteryl ester synthesizing enzymes also contribute to TAG synthesis, it remains to be established whether cholesteryl esters are capable of forming lipid droplets in the absence of triacylglycerols (Sandager et al., 2002). Here we show the existence of an alternative route for LD biogenesis, in which synthesis of another class of neutral lipids, retinyl esters, is sufficient to drive LD formation.

Based on biophysical properties of retinyl esters, we show that the spontaneous nucleation of retinyl ester-generated LDs is somewhat less efficient as compared to TAGs. Indeed, molecular dynamic simulations (Fig. 3 A,B) and *in vitro* experiments measuring the tension of artificial droplet interfaces (Table 1) suggest that RP droplets have a higher nucleation barrier, or lower nucleation efficiency than TAGs. Despite these biophysical considerations, retinyl ester-filled LDs are efficiently formed upon exogeneous induction of LRAT in yeast cells that lack the machinery to synthesize TAG/SE-filled LDs (Fig. 2). Surprisingly, the LRAT protein itself is involved in the generation of the large sized LDs that are characteristic for retinyl ester-containing LDs.

LRAT is broadly expressed with highest expression levels in the intestine, liver, testis and eye (Liu and Gudas, 2005; Ruiz et al., 1999). The majority of dietary vitamin A (∼75%) is, however, taken up by hepatocytes (Gottesman et al., 2001) and stored as retinyl esters in the liver, primarily in hepatic stellate cells (Blaner et al., 2009; Blomhoff et al., 1991; Friedman, 2008; Gottesman et al., 2001; Ross and Zolfaghari, 2004). LRAT expression is the highest in quiescent HSCs (Fig. 1E), in agreement with previous reports (Mederacke et al., 2013), but this expression is rapidly lost upon activation of HSCs. LRAT expression thus coincides with the presence of large LDs in quiescent HSCs. When LRAT expression is reduced (in activated HSCs, Fig. 1E) or absent in LRAT^−/−^ mice (Ajat et al., 2017; O’Byrne et al., 2005), HSCs lack their characteristic large LDs. These findings show that the presence of LRAT correlates with the existence of the large LDs, consistent with the data now presented here.

LRAT belongs to the papain-like or NlpC/P60 thiol protease superfamily of proteins (Anantharaman and Aravind, 2003). The H-RAS-like suppressor (HRASLS) enzymes, also known as LRAT-like proteins comprise a vertebrate subfamily that includes LRAT and that function in acyl chain remodeling of phospholipids (Golczak et al., 2012; Mardian et al., 2015). The *in vivo* biological substrates and activities of most family members have not yet been elucidated. LRAT is the best characterized family member, which catalyzes the formation of retinyl esters by transferring an acyl group from the *sn-l* position of PC onto ROH. The substrate specificity of LRAT for PC as acyl donor is intriguing. LD size is regulated by several factors including PC synthesis (Krahmer et al., 2011). During LD growth, the expanding LD monolayer activates CTP:Phosphocholine cytidylyltransferase, a key enzyme in PC synthesis. LRAT degrades PC which may result in activation of the same PC synthesizing pathway. Alternatively, reduced PC concentrations may either destabilize LDs and induce coalescence of LDs or increase ER bilayer tension, thereby generating larger (merged) LDs. Concomitant with retinyl ester production, however, PC is converted to lyso-PC. The bilayer surface tension is reduced by lyso-phospholipids, facilitating LD budding and reducing LD size (Ben M’barek et al., 2017). Thus, counteracting forces contributing to LD size are involved in the generation of retinyl ester-loaded LDs. The N-terminus of LRAT tips the balance as it is required for the generation of large LDs in mammalian cells. The effect of the N-terminus of LRAT on LD size is specific for retinyl ester-mediated LD formation: despite its presence in CHO-k1 LRAT-GFP cells, it does not affect the size of TAG filled LDs (Fig. 4). Since the N-terminus of LRAT does not seem to affect the rate of LD formation, it could change the thermodynamic equilibrium and decreases the nucleation concentration. As a result, it would then also dictate where LD formation occurs. However, we did not observe a colocalization of LRAT with nascent LDs (data not shown). Therefore, we consider it more likely that the observed specific affinity of the LRAT N-terminus for retinyl esters is likely to play an important role in affecting LD size, for example by changing the membrane topology of LRAT. LRAT is anchored to the ER membrane by a single transmembrane spanning domain at the C-terminus and with the catalytic domain and the N-terminus oriented towards the cytoplasm (Moise et al., 2007). The production of retinyl esters results in a strongly enhanced membrane binding of the N-terminus as illustrated by the deeper embedding of the N-terminus in the membrane, combined with the specific interaction with retinyl esters. Indeed, in the absence of the N-terminus, LRAT maintains retinyl ester production but fails to generate large LDs. In addition to a direct interaction with retinyl esters, it can however not be excluded that the N-terminus of LRAT also affects lipid droplet size by interfering with other mechanisms such as lipolysis and/or lipogenesis.

In summary, the molecular design of LRAT is well suited to store retinyl esters in LDs. With its C-terminal transmembrane domain, LRAT is anchored in the endoplasmic reticulum and directs newly synthesized retinyl esters in the lipid bilayer. The retinyl esters have the intrinsic capacity to form lipid droplets and with the N-terminal hydrophobic peptide LRAT specifically affects the size of retinyl ester-loaded lipid droplets.

## Acknowledgments

We thank Shreyas Sinha and Elmon Meijering for their assistance during the initial phase of experiments. Microscopy images were acquired at the Center of Cellular Imaging, Faculty of Veterinary Medicine, Utrecht University, we thank Esther van’t Veld for the technical assistance. Lipid analyses were performed at the Lipidomics Centre, Faculty of Veterinary Medicine, Utrecht University, we thank Jeroen Jansen for the technical assistance.

## Author Contributions

Conceptualization: J.B.H.; Methodology: M.R.M., T.A.W., A.R.T., W.A.P., J.B.H.; Software: T.A.W.; Formal Analysis: M.R.M., T.A.W.; Investigation: M.R.M., T.A.W., K.K.Y., A.T., M.C.M., L.C., A.C., M.W.H.; Writing – Original Draft: M.R.M., J.B.H.; Writing – Review & Editing: M.R.M., M.H., A.B.V., A.R.T., W.A.P., J.B.H.; Visualization: M.R.M., T.A.W., K.K.Y., M.C.M., L.C., A.C., R.W.W.; Supervision: M.H., A.B.V., F.R., A.R.T., W.A.P., J.B.H.; Project Administration: J.B.H.

## Declaration of Interests

The authors declare no competing interests.

## Material & Methods

### Hepatic stellate cells isolation and cell culturing

Hepatic stellate cells (HSCs) were isolated from 10–12 week-old male mice (C57BL/6J background, wild-type pubs from crossed LRAT^+/−^ heterozygote (Liu and Gudas, 2005 mice) as described before (Riccalton-Banks et al., 2003). Animals were handled according to governmental and international animal experimentation guidelines and laws. Experiments were approved by the Animal Experimentation Committee (Dierexperimentencommissie; DEC) of Utrecht University (DEC number 2013.III.02.016). After isolation, cells were protected from light and cultured on coverslips in 24-wells plates (Nunc, Roskilde, Denmark) in Dulbecco Modified Eagle Medium (DMEM) supplemented with 10% fetal bovine serum (FBS), 100 U/mL penicillin and 100 μg/mL streptomycin (all obtained from Gibco, Invitrogen GmbH, Lofer, Germany.

CHO-k1 cells were cultured in Ham’s F-12 medium supplemented with 7.5% FBS, 100 units/ml penicillin, and 100 μg/mL streptomycin. The human hepatic stellate cell line LX-2 cells (kindly donated by Dr. Friedman (New York, NY, USA)) were grown in DMEM containing 10% FBS, 100 U/mL penicillin and 100 μg/mL streptomycin. All cells were maintained in a humidified incubator (5% CO2) at 37 °C. Depending on the experiment, cells were incubated with retinol (Sigma, stocks of 30 mM in EtOH), oleic acid (Sigma) coupled to fatty acid free-BSA (Sigma, stocks of 10 mM fatty acid in 12% BSA) and/or triacsin C (Cayman, stocks of 1 mg/mL in DMSO).

### Retinol handling

Retinol (Sigma) was dissolved in ethanol (30 mM) and stored in 50 μL aliquots at −80°C. We routinely measured UV absorption spectra of retinol stocks before use and calculated the current stock concentration by making use of the molar extinction coefficient of retinol in EtOH (52,480 mol L^−1^ cm^−1^ at 325 nm) (Ross AC, 1981). In addition, we made use of the observation that after exposure to UV-light, a peak around 240 nm emerged, accompanied with a decrease around 325 nm. We routinely monitored E_325_/E_240_ ratios and used exclusively stocks with a ratio of 5 or higher, which was estimated to correspond with a retinol integrity of about 70%.

### Generation of stable cell lines

Human LRAT cDNA was synthesized and cloned into a pcDNA3.1(+) vector (Clontech, Palo Alto, California, USA) by a third party (GeneArt, Thermo Fisher Scientific, Waltham, MA, USA). In addition, the LRAT sequence was cloned into a pEGFP-N2 vector (Clontech), resulting in LRAT fused to the N-terminus of GFP. The construct coding for the deletion mutant (ΔNt-LRAT-GFP) was generated by site directed mutagenesis (New England Biolabs, Ipswich, MA). CHO-k1 and LX-2 cells were transiently transfected by Lipofectamine 2000 (Thermo Fisher) according to the manufacturer’s instructions. Stable CHO-k1 cells expressing the various LRAT variants or control GFP were generated as follows. Cells (2×10^6^) were electroporated in PBS with 15 μg DNA and a pulse of 260 V. After plating, cells were cultured for 48h in the absence and for 3 weeks in the presence of 1 mg/mL G-418 selection antibiotic (Thermo Fisher). Stable cells expressing GFP-fusion proteins were trypsinized and single GFP+cells with scatter properties (FSC and SSC) similar to their non-transfected counterparts were plated into 96-wells plates by FACS (Influx Cell Sorter, BD Biosciences). A GFP-negative gate was chosen to select monoclonal stable cells expressing non-fluorescent proteins. After plating, the monoclonal cells were allowed to grow until wells reached confluency. Clones with comparable morphology as compared to parental CHO-k1 cells (cell size, nucleus, lipid droplets) were selected. The absence or presence of LRAT enzymatic activity in LRAT, GFP, LRAT-GFP and ΔNt-LRAT-GFP expressing clones was confirmed by determination of retinyl esters (see below) and UV^+^-autofluorescent lipid droplets in combination with GFP-fluorescence by FACS (FACSCanto II, BD Biosciences) after incubation of cells with retinol.

### Confocal microscopy of mammalian cells

Cells were plated on Lab-Tek II 8-chamber slides (Thermo Fisher) and incubated as described in the figure legends. Cells were fixed in 4% paraformaldehyde (Electron Microscopy Sciences Hatfield, PA, USA). Subsequently, cells were stained with either DAPI, BODIPY 493/503 (Thermo Fisher), and/or LD540 (kindly donated by Dr. C. Thiele, Bonn, Germany) (Spandl et al., 2009). Staining for immunofluorescence was performed with anti-desmin or anti-α smooth muscle actin (both from Thermo Scientific), followed by goat-anti-mouse-alexa647 or donkey-a-rabbit-alexa647 (Life Technologies, Paisley, UK). Cells were mounted with FluorSave (Calbiochem, Billerica, MA, USA) and subsequently imaged with a Leica TCS SPE Laser Scanning Spectral Confocal Microscope (Wetzlar, Germany) or a Nikon A1R confocal microscope (Amsterdam, The Netherlands) using preset settings for the representative dyes. For the detection of retinoid autofluorescence, presets for DAPI were used.

### Quantification of cells with lipid droplet parameters

To quantify cells and lipid droplets, z-series of cells were imaged in tile-scan mode. Datasets were generated with either CellProfiler v2.2.0 (Kamentsky et al., 2011) or Imaris v8.2.0 (Bitplane, Belfast, Northern Ireland), both resulting in the identification of lipid droplets associated with individual cells. Data were subsequently expressed for individual cells containing ‘cell size’ (diameters of cells, in arbitrary units), ‘number of lipid droplets per cell’, ‘mean lipid droplet volume per cell’ (mean of all LD diameters of the cell to the 3^rd^ power, in arbitrary units) and ‘total lipid droplet volume per cell’ (product of ‘number of lipid droplets per cell’ and ‘mean lipid droplet volume per cell’, in arbitrary units). Subsequently, the resulting data were processed with R v3.4.4 and RStudio v1.0.153 using R-packages ‘openCyto’ v1.14.0, ‘reshape2’ v1.4.2 and ‘ggplot2’ v2.2.1.

To analyze the lipid droplet size distribution, lipid droplet diameters were not linked to individual cells, but analyzed per condition. Ranges of LD-diameter bins were calculated using the limits of the first, second, third, fourth and fifth diameter quintile of all lipid droplets. LD-presence in each bin was counted per condition and expressed as percentage (all lipid droplets per condition were set to 100%).

### Structured illumniation microscopy (3D-SIM)

After treatment, cells were stained with DAPI and HCS LipidTOX Red Neutral Lipid Stain (Thermo Fisher) and mounted with Vectashield Antifade Mounting Medium (Vector Laboratories, Burlingame, CA, USA). Structured illumniation microscopy was performed using a Deltavision OMX-V4 Blaze (GE Healthcare, Chicago, IL, United States) setup equipped with 4 sCMOS (PCO) cameras. Immersion oil with 1.516 refractive index (GE Healthcare) was placed on the 60x objective (Olympus U-PLAN APO, NA 1.42). Fluorophores were excited with a diode 405 nm (Vortran Stradus, 100 mW) and an OPSL 568 nm (Coherent, 100 mW) laser modulated to 1% by Neutral density filters. System supplied filter blocks were used to acquire fluorescence of DAPI (Ex: 382-409, Em: 421-450) and LipidTOX Red (Ex: 561-580, Em: 591-627). Raw images were processed using SoftWoRx software (GE Healthcare) with system OTFs pre-determined with 100 nm fluorescent polystyrene beads (Thermo scientific) and camera alignment parameters for the different channels (see supplemental materials). Acquired images were deconvolved using default settings (omitting Wiener filtering and background subtraction) including negative values (supplied as such in supplemental data) and intensities were linearly adjusted. Images in the figures are supplied as maximum intensity projections (movies are in supplemental data).

### Lecithin:retinol acyltransferase activity assay

Lecithin:retinol acyltransferase activity in homogenates expressing various LRAT variants was performed as described previously (Golczak et al., 2015). Briefly, cells were cultured overnight in T-75 culture flasks (CELLSTAR, Greiner Bio-One GmbH, Frick-enhausen, Germany) under normal cell growth conditions. After scraping in ice-cold PBS, cells were homogenized on ice with 26-gauge needles (BD Bioscience, San Jose, CA, USA). Homogenates containing 200 μg total protein were mixed with reaction mix containing 5 mM DTT, 5 mM EDTA, 10 mM Tris-HCl (pH 8.0), 1% BSA, 0.2 μM ascorbic acid, 2 mM PC(7:0/7:0) and increasing amounts of retinol. Subsequently, mixtures were incubated for 60 min at 37oC in amber glass vials. Levels of retinyl heptanoate were determined by LC-MS/MS (see below). Km en vmax values were estimated using Michaelis-Menten kinetics.

### Retinoids and neutral lipid determination by LC-MS/MS

Lipids were extracted as previously described (Bligh and Dyer, 1959). To avoid photo isomerization and oxidation of the retinoids, extractions were performed under red light and in amber tubes. In addition, 1 nmol butylated-hydroxytoluene was added to every sample. As internal standard, 250 pmol of retinyl acetate in MeOH/CHChl_3_ (1:1, v/v) was added. The combined chloroform phases were dried under nitrogen and stored at − 20 °C until further analysis.

Extract were dissolved in MeOH/CHCl3 (1:1) and stored in amber autosampler vials. To measure retinoids, samples were injected and separated on a 250 × 3.0 mm Synergi^™^ 4u Max-RP 80A column (4 μm particle size, Phenomenex, CO, USA) with a flow rate of 350 μL min^−1^. To this end, a gradient (solvent A; acetonitrile:water (95:5), solvent B; acetone:chloroform (85:15), 0 min; 90% A, 5 min; 40% A, 17 min; 0% A, 19 min; 90% A, 25 min; 90% A) was generated by a Flexar UHPLC system (Perkin Elmer, Waltham, MA, USA). The column outlet was connected to a triple quadrupole mass spectrometry (API 4000 QTRAP, MDS Sciex/Applied Biosystems, Foster City, Canada) with an atmospheric pressure chemical ionization (APCI) ionization source (set to 500 °C). Multiple reaction monitoring (MRM) in positive ion mode was used to detect retinyl ester species with settings and m/z transitions as described before (Ajat et al., 2017). Chromatographic peaks were integrated and quantified using Analyst software version 1.4.3 (Applied Biosystems, Foster City, Canada).

To measure other neutral lipids (sterols, triacylglycerols, steryl esters), samples were injected and separated on a Kinetex/HALO C8 column (2.6 μm, 150 × 3.00 mm; Phenomenex, Torrance, CA, USA). A gradient of methanol:H_2_O (5:5 v/v, solvent A) and methanol:isopropanol (8:2 v/v, solvent B) was generated by an Infinity II 1290 UPLC (Agilent, Santa Clara, CA, USA) and with a constant flow rate of 600 μL min^−1^ (0 min; 100% A, 2 min; 0% A, 8 min; 0% A, 8.5 min; 100% A, 10 min; 100% A). Lipids were measured using APCI in positive mode coupled to an Orbitrap Fusion mass spectrometer (Thermo Scientific, Waltham, MA, USA). Vendor data files were converted to mzML-format with msConvert (part of ProteoWizard v3.0.913) and processed with XCMS Online v3.7.0 (Tautenhahn et al., 2012).

### Growth and fluorescence microscopy of yeast

Yeast strains and plasmids used in this study are described in Supplemental Table 1. Yeast were grown at 30°C in synthetic complete media containing 0.67% yeast nitrogen base without amino acids (United States Biological), 2% glucose, and an amino acid mix (United States Biological). Retinol (Sigma-Aldrich) was added to the medium at 4 mM together with 1% Igepal CA-630 (Sigma-Aldrich). Cells were stained with 0.5 μg/ml BODIPY 493/503 (Invitrogen) for 10 min and being washed once with phosphate buffered saline.

Yeast were imaged at 30 °C in an Environmental Chamber with a DeltaVision Spectris (Applied Precision Ltd.) comprising a wide-field inverted epifluorescence microscope (IX70; Olympus), a 100 Å∼ NA 1.4 oil immersion objective (UPlanSAPO; Olympus), and a charge-coupled device Cool-Snap HQ camera (Photometrics).

### Western blot analysis

Cells expressing LRAT-GFP and ΔN-LRAT-GFP were growth to logarithmic growth phase and 10 OD600 units of cells were washed once with H_2_0 and lysed with glass beads in a Precellys 24 homogenizer (Bertin Instruments). The lysate was cleared by centrifugation at 500 *x g* for 10 minutes at 4°C. The protein concentration of the lysate was determined using a Bradford assay (Thermo Scientific) and 10 μg of protein were separated by 4–12% of SDS-PAGE gel (Invitrogen), and transferred to nitrocellulose membranes at 120 V for 2 h. The membrane was analyzed using primary antibodies to GFP (1:1,000; Roche) and anti-porin (1:1,000; Invitrogen). Proteins were visualized using IRDye secondary anti-mouse antibody (Li-COR Biosciences; 1:10,000). The blots were visualized with an Odyssey infrared imaging system (Li-COR Biosciences).

### LRAT mRNA expression by quantitative PCR

Expression of LRAT mRNA was determined as described (Tuohetahuntila et al., 2017). Briefly, RNA was isolated with a RNeasy Micro Kit (Qiagen, Venlo, The Netherlands) and cDNA was synthesized with an iScript cDNA Synthesis Kit (Bio-Rad). PCR amplifications were performed using a Bio-Rad detection system with iQ SYBR Green Supermix (Bio-Rad, Veenendaal, The Netherlands). Gene expression was normalized against reference genes, sequences of the primers are listed in Suppl. Table 2.

### Molecular dynamics

MD simulations were set up and run using the DAFT protocol (Wassenaar et al., 2015a), according to procedures for building membrane/solvent systems using INSANE (Wassenaar et al., 2015b) and for generating membrane/solvent/protein systems as described in (Wassenaar et al., 2015a). The latter comprises a step for the generation of the LRAT-Nt peptide (sequence MKNPMLEVVSLLLEKLLLISNFTLFSSGAAGEDKGRNSF; secondary structure LLLLHHHHHHHHHHHHHHHLLLLLLLLLLLLLLLLLLLL) or the CCTα P2 peptide (sequence VEEKSIDLIQKWEEKSREFIGSFLEMFG; secondary structure HHHHHHHHHHHHHHHHHHHHHHHHHHHH), which was placed horizontally above the membrane. For docking simulations, the helix was placed at a distance 4 nm from the bilayer center with random rotation around the helix axes, while for assessment of the effect on lens formation speed, the helix was placed at 2.5 nm from the bilayer center, with the hydrophobic side facing towards the membrane. Simulations were performed using the coarse grain Martini 2.2 model (Marrink et al., 2004, 2007; Monticelli et al., 2008).

Simulations for assessing binding of LRAT-Nt AH to membranes with different compositions were set up in a hexagonal prism unit cell with a total number of 250 lipids per leaflet and a height of 9 nm. Simulations for assessing the lens forming propensity and speed, were set up in a hexagonal prism unit cell with base length of 24 nm and height of 10 nm, regardless of the presence or absence of protein. Numbers or ratios or lipids were set as described in the main text. All simulations were performed using Gromacs 2018.x (Pronk et al., 2013), using the automated Martini workflow *martinate* (Wassenaar et al., 2013).

### Electron Microscopy

Yeast cells were grown in synthetic drop-out media lacking leucine at 30°C to an OD600 of approximately 0.6. IGEPAL (Sigma-Aldrich) was added to the cell cultures to a final concentration of 1%, prior to addition of retinol (Sigma-Aldrich) to a final concentration of 2 mM. Cells were then incubated for 10 min at 30°C before being processed for electron microscopy as follow. Cells were chemically fixed, embedded with 12% gelatin, cryo-sectioned and stained as previously described (Griffith et al., 2008). Sections were imaged in a FEI CM100bio electron microscope at 80 KV, equipped with a digital camera (Morada; Olympus). Two different grids with sections obtained from the same preparation were statistically evaluated by counting 75 randomly selected cell profiles before determining the average number of lipid droplets per cell section plus standard deviation between the two grids.

### Monolayer tension measurements

Tension measurements were performed using a drop tensiometer device (Tracker, Teclis-IT Concept, France) (Ben M’barek et al., 2017). The principle of the drop profile analysis is based on the determination of the shape of a liquid drop suspended in another liquid form from a video image and its comparison with theoretical profiles calculated from the Gauss Laplace equation. The retinyl palmitate (Sigma, R-1512) drop (neutral lipid phase), with or without containing the DOPC phospholipid, was formed in buffer (50mM HEPES, 120mM potassium acetate, 1mM magnesium chloride, pH 7.4) at room temperature. The tension was allowed to stabilize for a few minutes (it decreases by the continuous absorption of phospholipids to the oil/water interface). Then, the drop is compressed by decreasing its volume, until complete saturation of the interface is reached (marked by a plateau of tension during compression).

### Droplet interface bilayer (DIB) experiments

The DIB experiments was performed following the previous study of (Ben M’barek et al., 2017). An oil phase containing phospholipids was prepared first. DOPC, DOPE and DOPA were purchased from Avanti Polar Lipids, Inc. Lipids were mixed to the RP (Sigma, R-1512) oil at a final lipid concentration of 0.2% w/w (about 10% chloroform was in the final mixture in the case of DOPC initial stabilization of the DIBs, and we let it evaporate over time; for the other phospholipids, chloroform was evaporated prior to addition of RP). Then, an emulsion was prepared mixing the buffer with RP (1:5 v/v). Finally, the same volumes of emulsion and phospholipids in RP were put together, and the resulting emulsion was placed on a hydrophobic coverslip.

### Interfacial tension measurements

A pendent droplet tensiometer designed by Teclis Instruments (Longessaigne, France) was used to measure the interfacial tension of oil/water interfaces. All experiments were conducted at room temperature. To create oil/buffer interfaces, oil drops (10μL) were formed at the tip of a J-needle submerged in 5 mL of HKM buffer.

#### Oil-peptide surface tension measurement

10 μl of LRAT peptide solution in DMSO was added to the bulk buffer while the tension of trioleoylglycerol/buffer or retinyl palmitate/buffer interface was measured. The resulting concentration of peptide was 4.6 μM for both retinyl palmitate and trioleoylglycerol experiments. Oil-peptide surface tension was determined when the interfacial tension was stabilized.

After tension stabilization, the interface is compressed. The compression was then stopped, and area was kept constant as interfacial tension was recorded. Relaxation of the tension was observed only for trioleoylglycerol/buffer interfaces. Human LRAT peptide (AA 1-39) was synthesized by Bio-Synthesis (Lewisville, Texas, USA).

### Statistical analyses

All figures were built in RStudio v1.0.153 (R v3.4.4) and processed in Inkscape v0.92.2. Barplots represent means ± standard deviation (SD) or standard error of the mean (SEM) as indicated in the figure legends. Statistical significance was determined by two-tailed paired or unpaired Welch’s t-test or Wilcoxon rank sum test, or by Pearson’s Chi-squared test, as indicated. In experiments with multiple testing, P-values were corrected by the Benjamini-Hochberg procedure. P-values below 0.05 were considered statistically significant.

