## Supplemental Information for "Lecithin:Retinol Acyl Transferase (LRAT) induces the formation of lipid droplets"

Molenaar *et al.*

### Supplemental Figures

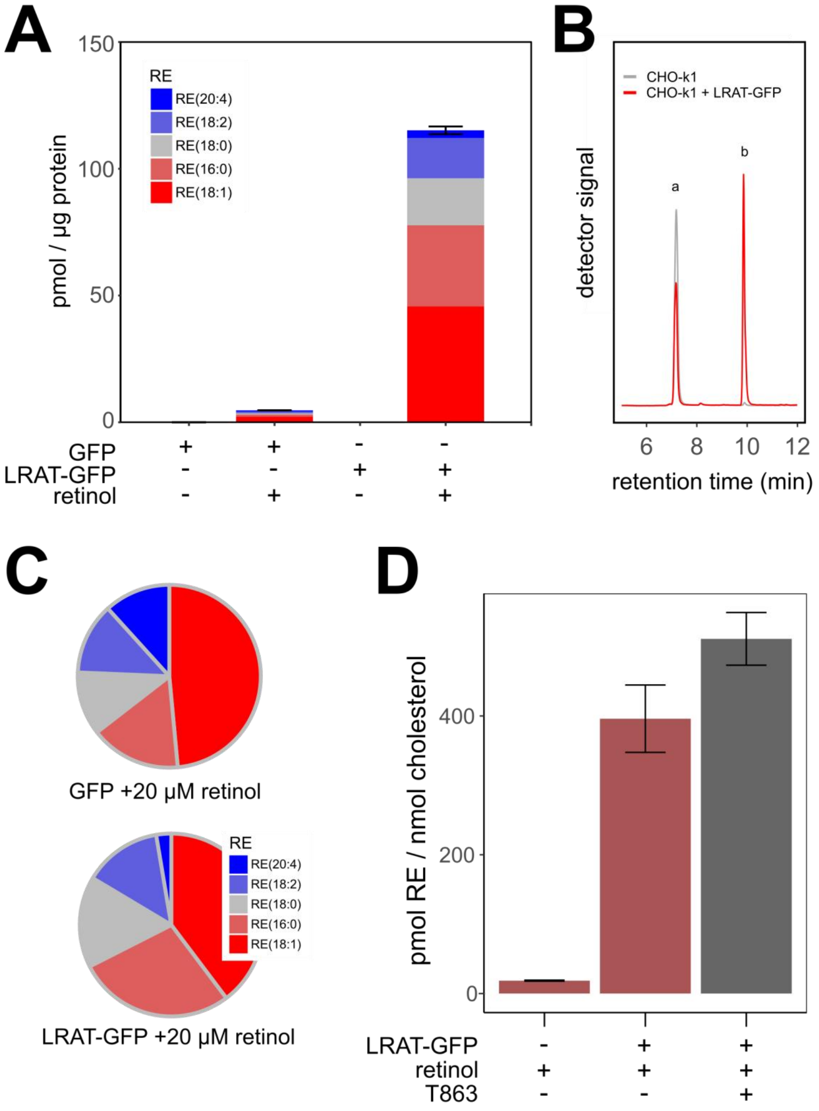


**Figure S1. Quantification of retinyl ester content in CHO-k1 cells. (A)** CHO-k1 cell lines expressing GFP or LRAT-GFP were incubated in the presence or absence of 20 µM ROH. RE species were analyzed by LC-MS/MS, amounts are expressed as pmol RE per µg protein (means ± SD of a representative experiment. **(B)** Chromatogram showing *in vitro* LRAT-activity of CHO-k1 homogenates with (red line) and without (gray line) expressing LRAT-GFP. After incubation with PC(7:0/7:0) and 10 µM ROH, ROH (peak a) and RE(7:0) (peak b) were measured by LC-MS/MS. MRM-transition 269/93 (retinoid backbones) is shown. **(C)** Relative contribution of retinyl ester species that were synthesized under the conditions as described in panel A. **(D)** CHO-k1 cell lines with or without expression of LRAT-GFP were incubated overnight with 20 µM ROH in the presence or absence of 10 µM DGAT1-inhibitor T863. Total RE (m/z 269) and free cholesterol (m/z 369) were analyzed by LC-MS. Amounts are expressed as pmol RE per nmol cholesterol (means ± SD of a representative experiment).

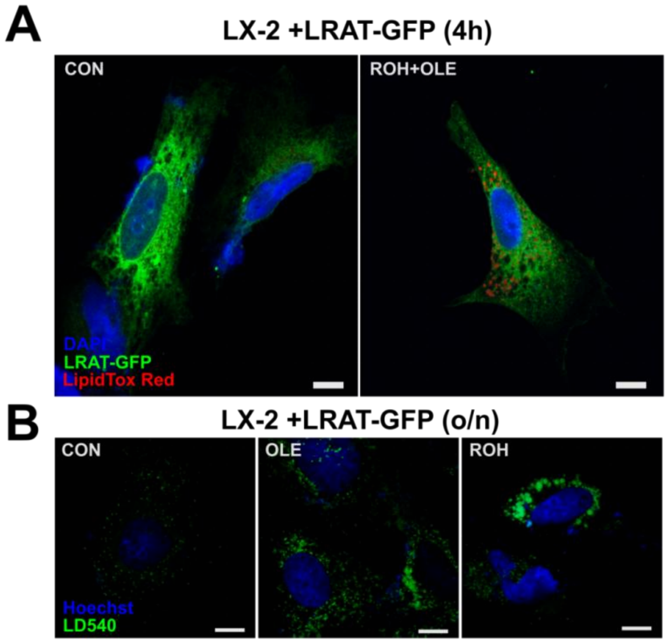


**Figure S2. LRAT and lipid droplet localization in LX-2 cells. (A)** Confocal microscopy of transiently transfected LX-2 cells expressing LRAT-GFP (green), co-stained with DAPI (blue) and LipidTox Red (red). Cells were incubated without (left) or with (right) 20 µM ROH and 200 µM OLE for 4 hours. **(B)** Confocal microscopy of transiently transfected LX-2 cells expressing LRAT-GFP, stained with DAPI (blue) and LD540 (green). Cells were incubated without (left) or with 200 µM OLE (middle panel) or 20 µM ROH (right panel) overnight. Scale bars indicate 10 µm.

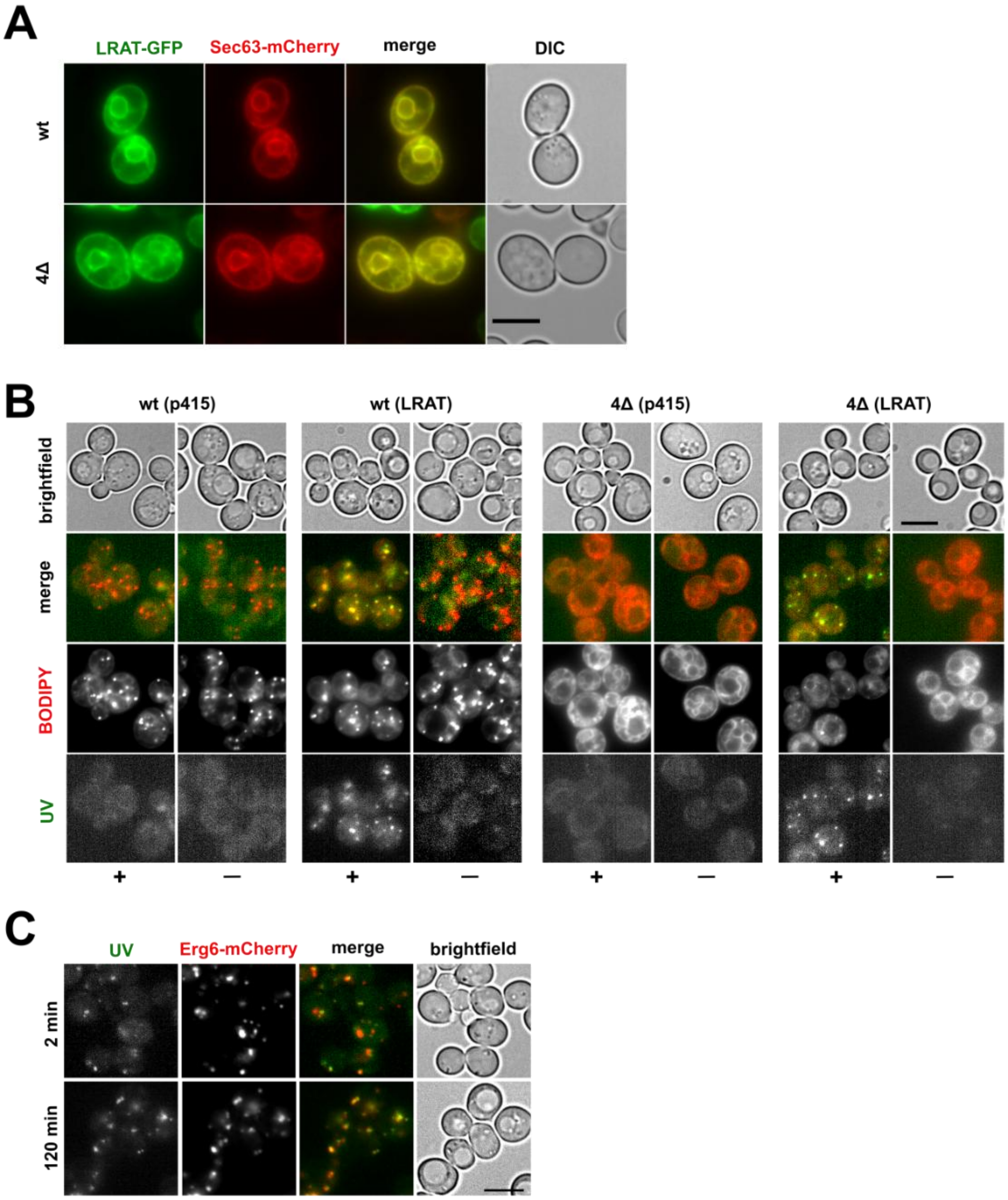


**Figure S3. LRAT-mediated LD-formation in the absence of pre-existing LDs. (A)** Wide-field microscopy of *wt* and 4Δ yeast cells, expressing LRAT-GFP (green) and Sec63-mCherry (red), showing DIC (white) and fluorescence channels (colors and merge). **(B)** Wide-field microscopy of *wt* and Δ4 yeast cells, with or without expressing LRAT, 2 hours after incubation with or without 2 mM ROH ('—' or '+'). After staining, images of UV-autofluorescence (green), BODIPY (red) and brightfield were taken. **(C)** Wide-field microscopy *wt* yeast cells, expressing LRAT and Erg6-mCherry (red). Cells were imaged 2 (top) or 120 (bottom) minutes after addition of 2 mM ROH. Images of UV-autofluorescence (green), BODIPY (red) and brightfield (white) were taken. Scale bars indicate 5 µm.

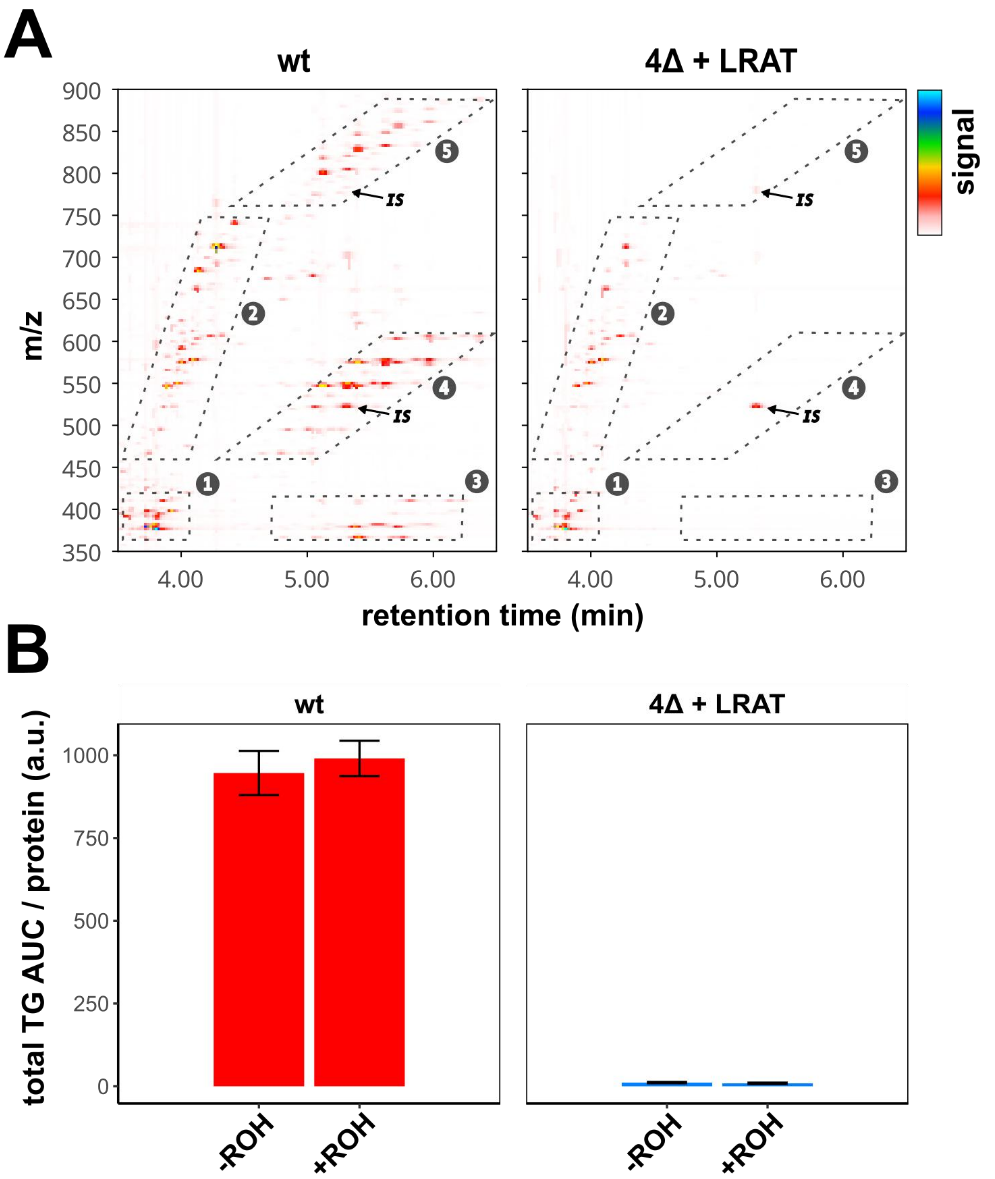


**Figure S4. Relative quantification of triacylglycerols in yeast by LC-MS. (A)** LC-MS contour plots of retention times between 3.5 and 6.5 min (x-axis) and m/z-values between 350 and 900 (y-axis). Detector signal is represented by color code (see legend). Numbered boxes indicate areas with ions from (1) sterols, (2) diacylglycerols and ceramides, (3) steryl esters and triacylglycerols $[M+H-2RCOOH]^+$, (4) triacylglycerols $[M+H-RCOOH]^+$ and (5) triacylglycerols $[M+H]^+$. IS: internal standard TG(15:0/15:0/15:0). Representative samples of wild-type (left) and 4Δ+LRAT (right) yeast strains, incubated with 2 mM ROH, are shown (for experimental details, see Fig. 2). **(B)** Relative quantification of summed peak area of all triacylglycerols ions except internal standard from box 5 (panel A) of *wt* (left panel, red) and Δ4+LRAT (right panel, marine) yeast cells, in the presence or absence of 2 mM ROH. Barplot indicates means ± SD.

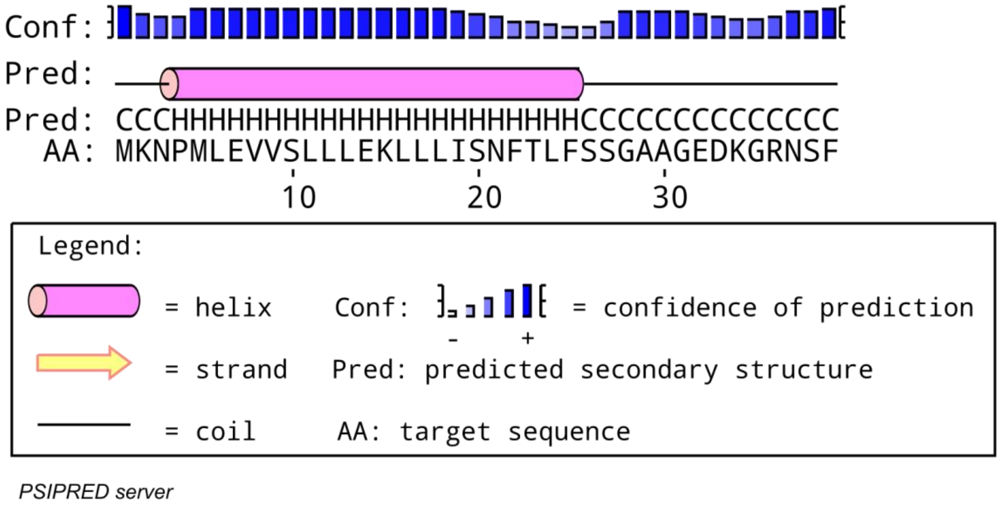


**Figure S5. PSIPRED secondary structure prediction of LRAT N-terminus.** Secondary structure prediction of N-terminus of LRAT by the PSIPRED-server (Buchan et al., 2013). Predicted secondary structures are ndicated by pink cylinders (alpha-helices, ‘H’), yellow arrows (β-stands, ‘S’) and black lines (coils, ‘C’). Blue bars indicate the prediction’s degree of confidence.

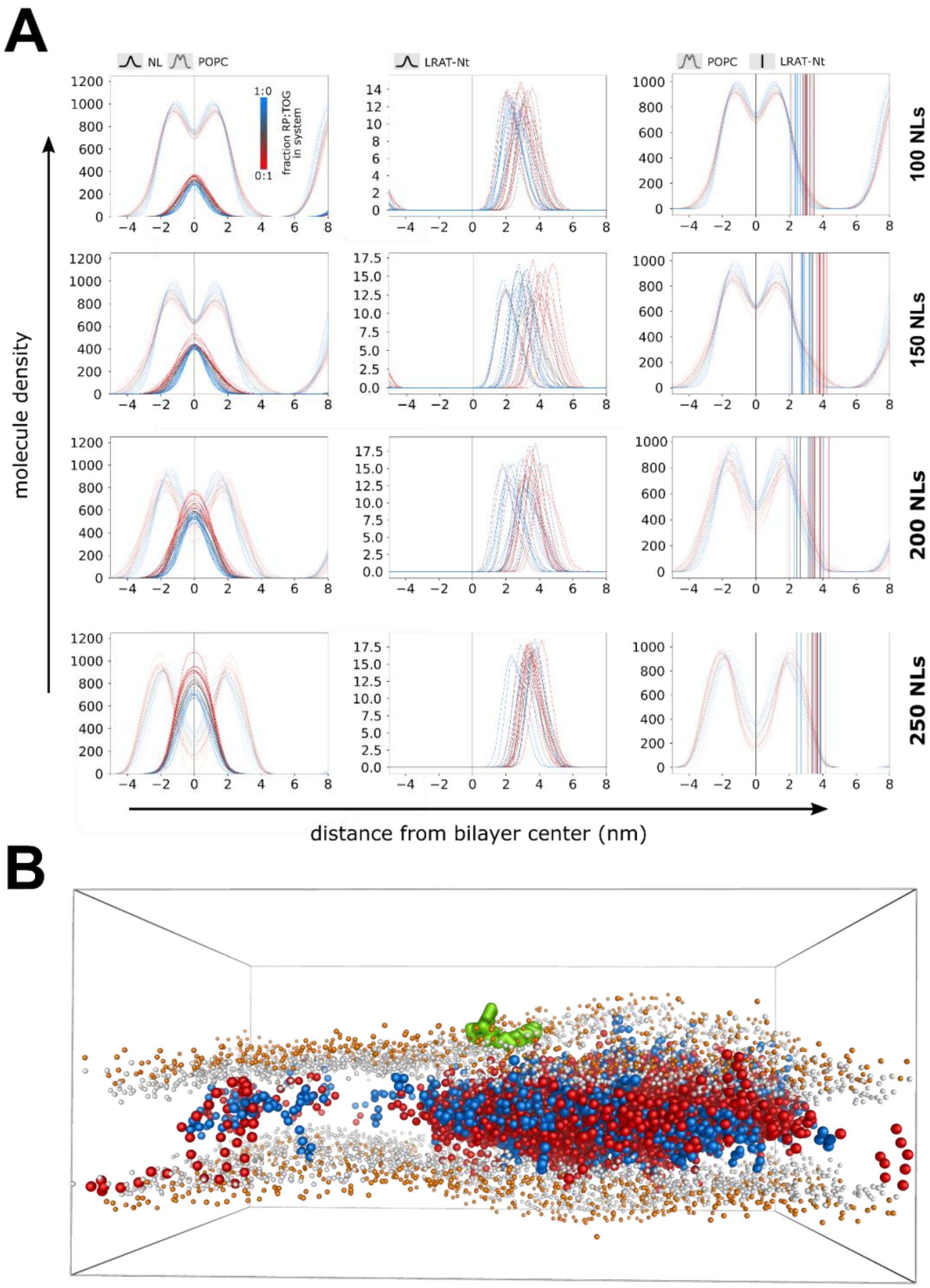


**Figure S6. Binding of Nt-LRAT to LD lenses**. **(A)** Molecule densities as function of distance from bilayer center of neutral lipids (bright lines, upper panels), POPC (opaque lines, upper and bottom panels) and LRAT-Nt (middle panels). Colors are scaled to neutral lipid compositions in the simulations (marine: pure RP to red: pure TOG). The vertical lines in the bottom panels indicate the positions Nt-LRAT's centers of mass. **(B)** Representative side view of a simulation endpoint illustrating the affinity of LRAT-Nt for RP. Neutral lipids are shown as large beads (marine: RP, red: TOG), POPC molecules as small beads (grey: glycerol-groups, orange: phosphate-groups) and the LRAT-Nt peptide in green.

### Supplemental Tables

**Table S1. S. cerevisiae stains and plasmids used in this study.**

| strain/plasmid | genotype or description | source |
|---|---|---|
| BY4742 | Mat α *his3*Δ1 *leu*2Δ0 *lys*2Δ5 *ura*3Δ0 | laboratory collection |
| VCY02 | BY4742 MATα *his3Δ1 leu2Δ0 lys2Δ0 ura3Δ0 met15Δ0, are1::KanMX6, are2Δ::KanMX6, trp1::URA, lro1::TRP1, dga1::lox-HIS-lox* | Jacquier *et al.* 2011 |
| VCY63 | BY4742 Mat α *his3Δ1 leu2Δ0 lys2Δ5 ura3Δ0 Erg6-mCherry::HIS3* | this study |
| VCY64 | BY4742 *MATα his3Δ1 leu2Δ0 lys2Δ0 ura3Δ0 met15Δ0, are1::KanMX, are2Δ::KanMX, trp1::URA, lro1::TRP1dga1::lox-HIS-lox Erg6-mCherry::HIS3* | this study |
| KKE018 | p415GPD-LRAT *(LEU2/CEN*), full length LRAT express under *GPD1* promoter | this study |
| KKE020 | p415GPD-LRAT-GFP *(LEU2/CEN*), full length LRAT-GFP express under *GPD1* promoter | this study |
| NOY53 | MATa *his3Δ1 leu2Δ0 met15Δ0 ura3Δ0 scs3::HIS3 yft2::KanMX* | Choudhary *et al.*, 2015 |
| pAT124 | YCplac33-*SEC63*-mCherry (*URA3/CEN*), Sec63-mCherry expressed under *SEC63* promoter | this study |
| pcDNA3-LRAT | full length human LRAT | this study |
| pEGFP-N2-LRAT | full length human LRAT fused to EGFP | this study |
| pEGFP-N2-ΔNt-LRAT-GFP | N-terminal deletion (Δ1-36) of human LRAT fused to EGFP | this study |

**Table S2. Quantitative PCR primers used in this study.**

| gene | species | | source | | |
|---|---|---|---|---|---|
| Lrat | Mus musculus | | Tuohetahuntila *et al.*, 2017 | | |
| Ywhaz | Mus musculus | | Tuohetahuntila *et al.*, 2017 | | |
| Hmbs | Mus musculus | | Tuohetahuntila *et al.*, 2017 | | |
| Hprt | Mus musculus | | Tuohetahuntila *et al.*, 2017 | | |
| Gapdh | Mus musculus | | this study | | |
| | | **primer** | | **5'-sequence-3'** | **$T_m$ (°C)** |
| | | F | | GAA GGT CGG TGT GAA CGG | 61 |
| | | R | | TGA AGG GGT CGT TGA TGG | |
| Actb | Mus musculus | | this study | | |
| | | **primer** | | **5'-sequence-3'** | **$T_m$ (°C)** |
| | | F | | AGC TCC TTC GTT GCC GGT CCA | 57 |
| | | R | | TTT GCA CAT GCC GGA GCC GTT G | |
| Rps18 | Mus musculus | | this study | | |
| | | **primer** | | **5'-sequence-3'** | **$T_m$ (°C)** |
| | | F | | GAT CCC TGA GAA GTT CCA GCA C | 57 |
| | | R | | ACC ACA TGA GCA TAT CTC CGC | |

### Supplemental Videos

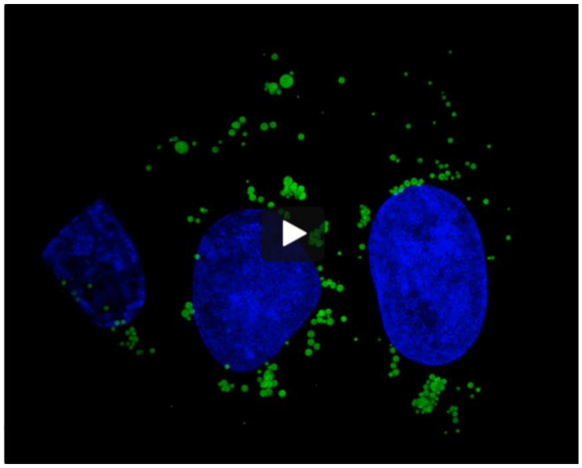

Video S1.

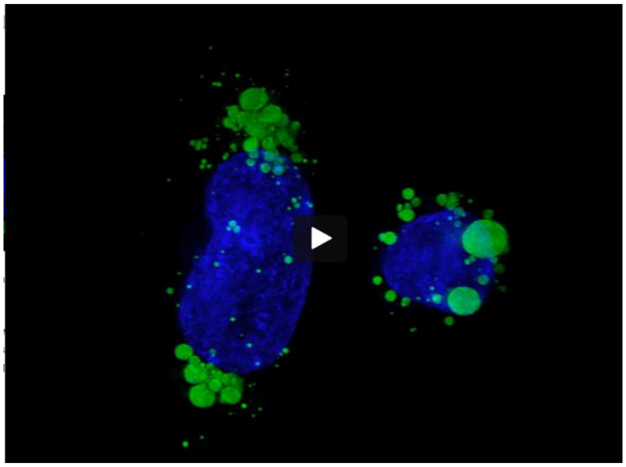

Video S2.

**Video S1-2. 3D reconstructions of LDs in LRAT-GFP.** 3D reconstructions by SIM of LRAT-GFP expressing CHO-k1 cells incubated with 200 μM OA **(Video S1)** or 20 μM ROH **(Video S2),** showing DAPI (blue) and LipidTox Red (green).
